# Digital twins forecast individual immune trajectories

**DOI:** 10.64898/2026.10.23.761347

**Authors:** Chengpeng Hu, Chenpeng Yu, Calvin Yu-Chian Chen

## Abstract

The human immune system differs markedly between individuals and changes in response to infection, vaccination and treatment^1–3^. AI virtual cells are being built to simulate cellular responses to perturbation ^4–6^. Yet these models, like most immune-response predictors, describe each person by a single pre-challenge measurement^7–11^ and forecast change from patterns pooled across people. Digital twins, updated with every new observation of an individual, could forecast how that person’s immune state changes ^4,12^. Here we present ImmuneTwins, which builds these twins from single-cell profiles by personalizing a shared virtual-cell model with each individual’s observations. In two external populations, ImmuneTwins reduced immune-age error by 32% and 42% relative to the best existing method. During repeated influenza vaccination, its twins predicted the direction of immune-cell change 35% more accurately than the best published method, and their error fell by 27% as observations accumulated. The twins revealed that immune cells and plasma proteins recover about half their deviation from a persistent personal reference within 90 days, whereas response size varies between challenges. They further localized B-cell remodelling in exceptional longevity and anticipated patient-specific immune trajectories during myeloma treatment. These results establish virtual-cell digital twins for forecasting individual immune change and identifying each person’s characteristic immune features.

## Introduction

The immune system protects against infection and maintains tissue homeostasis through the coordinated activity of diverse cell populations ^13^. Genetic background, microbial encounters and environmental exposures shape these populations through-out life, producing substantial differences in immune composition and function between people ^1,14–18^. Large-scale profiling has linked immune variation to infection susceptibility ^19^ and yielded measures of immune age and health ^20–22^ that place each person within population variation and are associated with disease burden, survival and treatment outcomes ^23–26^. These measures, however, are usually developed within particular cohorts ^26^, and their estimates can shift systematically when applied to populations that differ in age distribution or measurement conditions ^27^. Group-level comparisons can also obscure differences between individuals of the same age or diagnosis. In monogenic immune disorders, inter-individual differences account for more variance in many immune measurements than disease category does ^25^. Age or diagnosis alone therefore provides an incomplete account of a person’s immune composition and function. Models of individual immune state need to capture these differences and remain accurate when applied across populations.

Single-cell atlases allow these differences to be examined across immune lineages, tissues and human populations ^28–31^, providing the data on which virtual cells simulate cell states and their responses to perturbation ^4,32,33^. Multimodal integration and foundation models learn universal representations of cell states from large reference datasets ^34–37^. Models such as PaSCient and PULSAR combine these cell-level representations to characterize patient phenotypes ^11,38,39^. For the immune system, virtual cells need to resolve these cell states: gene expression within a cell population can change even when its proportion remains stable ^2,40^, and such changes are lost when a sample is reduced to major cell-type proportions or averages across all cells ^41,42^. They also need to generalize beyond the contexts in which they were trained ^6,43^. A virtual cell trained in one cohort and applied to another population encounters differences in cellular distributions and study conditions ^27^, which can bias its individual-level predictions ^44^ and make it unclear whether a difference reflects immune state or cohort-specific effects ^45^. Comparisons across populations therefore require virtual cells that preserve individual biological differences despite these changes in study context.

Following the same person over time reveals which immune features remain stable and which change after immune challenge ^2,46^. Previous exposures can leave lasting effects ^47,48^, and measurements made before a challenge can help predict the subsequent response ^7–9,40^. Predicting such responses is a central task of virtual cells ^4,33^: cellular perturbation models and multicellular foundation models predict how immune cells and individuals respond to an intervention ^5,10,11,49,50^. Yet these predictions describe each person by a single measurement and forecast change from patterns pooled across people. The individual’s own earlier observations, which longitudinal studies show to carry stable and lasting features, are not used to anticipate how that person will change. Modelling each person from their own record of observations is the premise of digital twins.

Digital twins build such individual models and update them as new observations are made ^12^. Organ digital twins reproduce the physiology of individual ex vivo human lungs to evaluate therapies ^51^, and longitudinal clinical models use each person’s recorded history to forecast disease trajectories and complete missing measurements ^52,53^. A proposed next step is to combine structure shared across patients with each person’s own trajectory, distinguishing stable personal baselines from individual change ^53^. The AI virtual cell roadmap envisions such models at cellular resolution: personalized virtual cells, built from a patient’s blood single-cell profiles and refined with each new measurement, that anticipate changes in health and the outcomes of treatment^4^. Single-cell digital twins have been generated from each blood sample separately, with real-time immune monitoring left as a future direction ^54^. The immune system, accessible through repeated blood sampling, is a natural candidate for this approach. Immune states differ persistently between people and change transiently with each challenge, so a twin must learn how immune cells respond across people while remaining specific to each individual, at the resolution of the cell states in which these differences arise. Such a twin could forecast how a person’s immune state will change and reveal which of its features are characteristic of that person.

Here we present ImmuneTwins and demonstrate that a virtual-cell model, personalized with each individual’s own record of immune responses and resting states, can act as a digital twin of the human immune system, forecasting from blood single-cell profiles how a person’s immune cells will change after vaccination and during cancer treatment. ImmuneTwins generates each person’s future immune cells from their current cells with one virtual-cell model shared across people, and updates each twin with every new visit. Its patient representation, learned from 417 participants of the Chinese Immune Multi-Omics Atlas, predicted immune age zero-shot in 1,215 participants from two external cohorts of different ancestry. In 75 participants followed through two annual influenza vaccinations and 20 patients receiving myeloma induction therapy, the twins forecast individual immune-cell changes more accurately than published perturbation and representation models, and their vaccination forecasts improved with each additional visit. The twins revealed that each person’s immune system has its own resting state to which it returns: vaccination pushed it away by a different amount each year, but immune cells and plasma proteins recovered about half of their deviation from this personal reference within 90 days. In silico exchange of cells between individuals traced the stronger secretory programme of supercentenarians to memory and plasma-like B-cell states within a smaller B-cell compartment, and in myeloma, each patient’s early treatment response anticipated later host immune changes, including cytotoxic-programme decline and myeloid remodelling, beyond those reflected in tumour response.

## Results

ImmuneTwins builds a personal digital twin for each individual on a shared base model, a virtual-cell model that learns a patient-level representation of immune-cell states and generates an individual’s future cells from their current cells (Fig. 1). Each twin adds two personal components to the base model: personal history, the responses the individual has already completed, and a personal reference, the individual’s usual composition estimated from earlier resting visits. We evaluated whether the shared representation generalized across populations, tested whether personal observations distinguished the future cell states of individuals after vaccination, identified the persistent individual feature on which these forecasts rely and applied the twins to exceptional longevity and myeloma treatment (Supplementary Figs. 1 and 2).

**Fig. 1.**
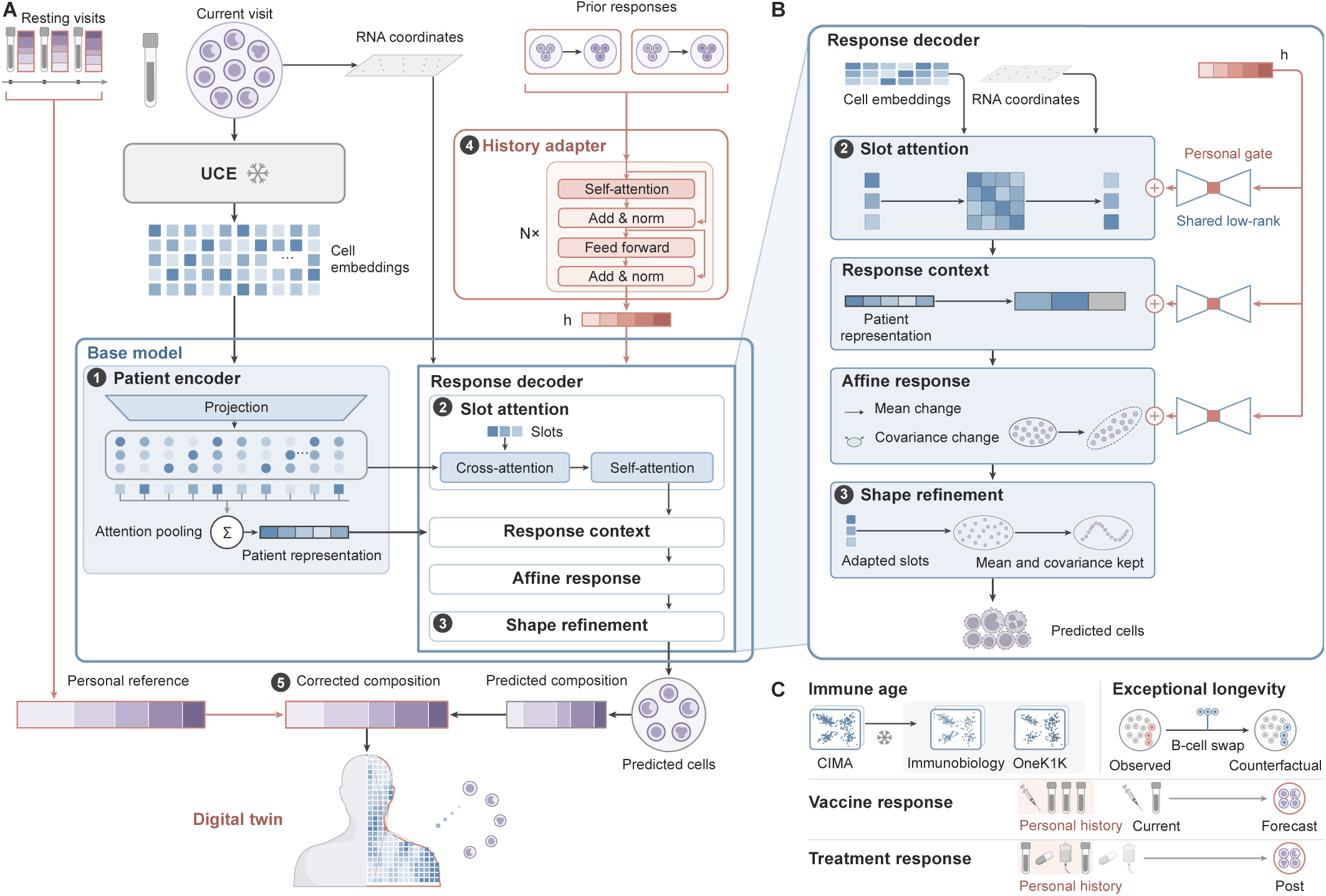
ImmuneTwins builds personal digital twins of the immune system by adapting a shared single-cell model to each individual’s own observations. **A**, Cells from the current visit are embedded by the frozen Universal Cell Embeddings (UCE) model and pooled by the patient encoder (1) into a patient representation, from which the response decoder (2, 3) predicts the cells at a later visit. The encoder and decoder form the base model shared by all individuals. A digital twin adds two personal components: a history adapter (4) that encodes the individual’s prior responses into a history embedding h, which adapts the decoder, and a correction of the predicted composition towards the personal reference, the individual’s usual composition across earlier resting visits (5). **B**, The response decoder. Slots attend to cell embeddings and RNA coordinates (2) and are combined with the patient representation into a response context, which sets an affine change in the mean and covariance of the cell distribution; shape refinement then reshapes the distribution while keeping its mean and covariance (3). Through personal gates on shared low-rank maps, h adapts these steps. **C**, Applications: zero-shot transfer of immune age from the Chinese Immune Multi-Omics Atlas (CIMA) to the Immunobiology of Aging and OneK1K cohorts, in silico exchange of B cells between supercentenarians and controls, and forecasts of individual responses to influenza vaccination and myeloma treatment from personal history. Snowflakes mark frozen models.

**Fig. 2.**
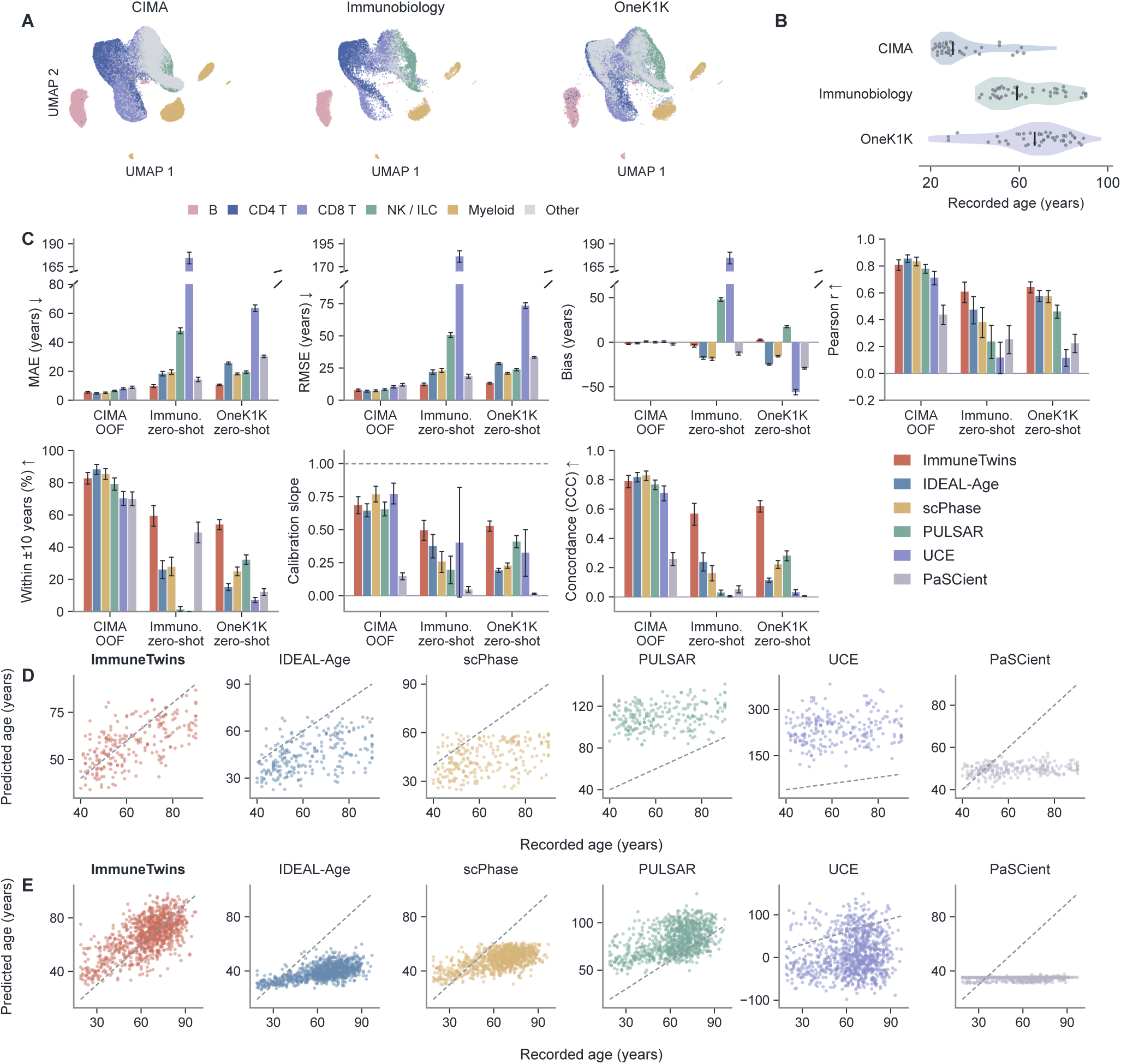
Cross-population transfer of immune-age predictions. **A**, UMAP of cell representations from the CIMA, Immunobiology and OneK1K cohorts. **B**, Age distributions of the three cohorts. **C**, Age-prediction performance of six methods within CIMA and in the two external cohorts. **D,E**, Predicted versus recorded age for each method in Immunobiology (D) and OneK1K (E).

### Cross-population immune-age prediction

ImmuneTwins learned patient-level aggregation and age prediction from 417 participants in the Chinese Immune Multi-Omics Atlas (CIMA) ^31^. To test whether this representation generalized beyond the context in which it was learned, the fitted model was applied zero-shot to the Immunobiology of Aging cohort (hereafter Immunobiology) ^2,55^ and to the Northern European ancestry OneK1K cohort ^29^, which differ from CIMA in ancestry, study source and age distribution (Fig. 2A,B).

Within CIMA, ImmuneTwins performed comparably to the strongest of the five comparator methods ^11,27,37,38,56^ (Fig. 2C). Cross-population evaluation, however, exposed systematic shifts in the age predictions of all five comparators, whose mean predictions departed from recorded age by more than 12 years in both external cohorts. These offsets included both overestimation and underestimation, and several methods also compressed participants of different ages into narrow prediction ranges, with calibration slopes as low as 0.01 (Fig. 2C–E). An offset of this size places an individual more than a decade from their recorded age before any biological difference is considered, limiting the comparability of immune-age estimates across populations. In both external cohorts, ImmuneTwins outperformed all five comparators on every metric: it reduced the mean absolute error to 9.7 and 10.6 years, 32% and 42% below the best comparator in each cohort, estimated 59% and 54% of participants within ten years of their recorded age, compared with at most 49% and 32%, and limited the systematic offset to *−*4.0 and +2.7 years (Fig. 2C–E). Learned aggregation was particularly important for this transfer: replacing it with mean pooling increased external errors more than errors within CIMA (Supplementary Fig. 3). The representation underlying the virtual-cell model thus generalized zero-shot to new populations and serves as the base model of every twin.

### Forecasting individual vaccine responses

We next asked whether personal observations could distinguish how individuals with similar current immune states would respond to vaccination. In 75 held-out participants of the Sound Life cohort undergoing repeated influenza vaccination ^2^, each twin generated virtual cells for the second-year day-90 visit from the cells of the pre-vaccination visit, using the completed first-year response and earlier resting visits as personal observations.

ImmuneTwins achieved the best overall performance among the methods compared, which included the base model without personal components, published perturbation and representation models ^10,11,37,49,57^, and forecasts assuming no change or the population-average change (Fig. 3A). Its advantage was greatest in the direction of change: the cosine similarity between predicted and measured mean changes reached 0.37, 35% higher than the 0.27 of the best published method, trVAE, and 41% higher than the base model, and ImmuneTwins outperformed every method that predicts a change (*q <* 0.05). ImmuneTwins also ranked first for within-season forecasts from day 7 to day 90 (Supplementary Fig. 4).

**Fig. 3.**
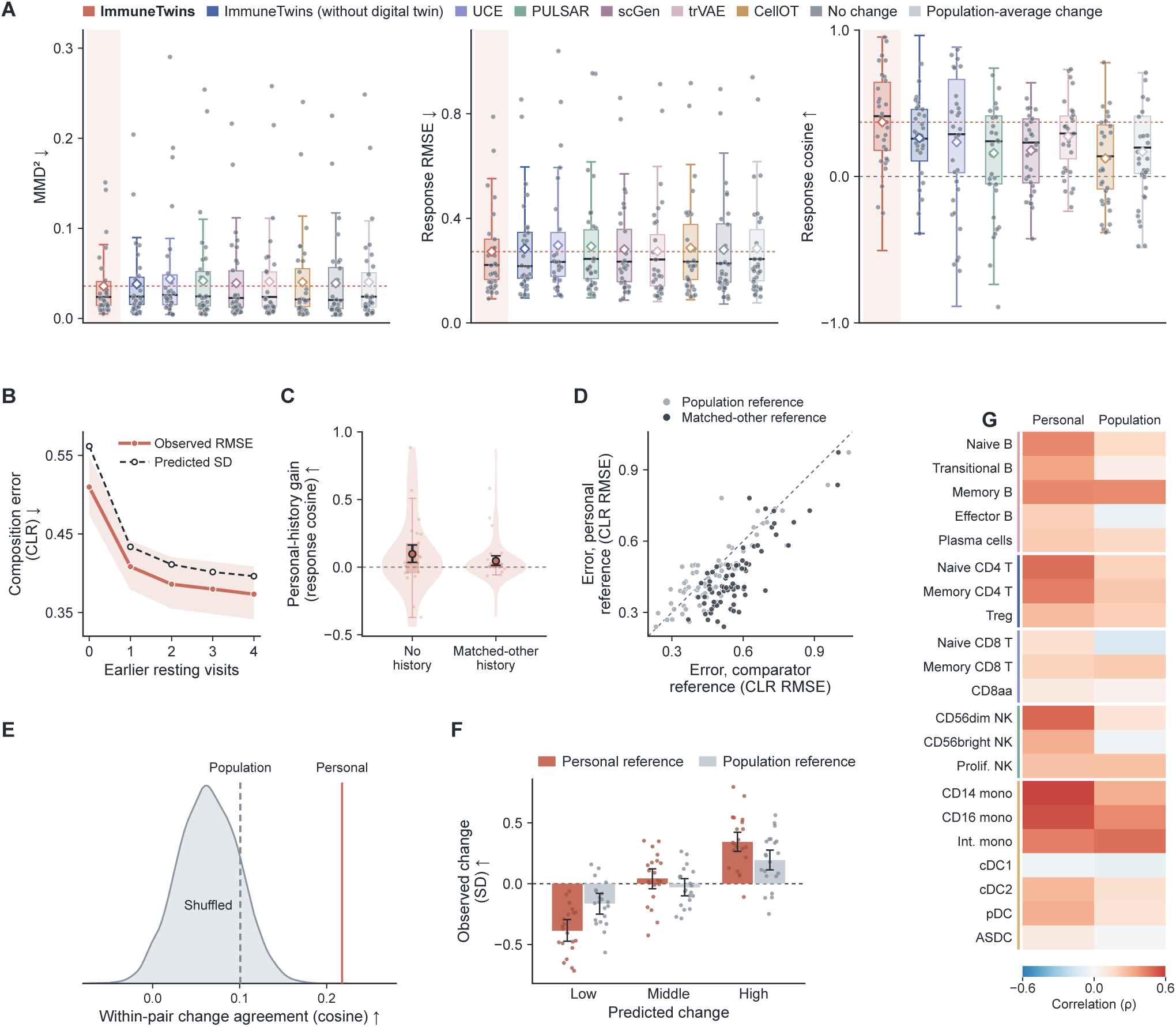
Personal immune-state forecasts improve with longitudinal observations. **A**, Second-year day-0-to-day-90 forecasts by ImmuneTwins with and without the digital twin, published methods and simple baselines. **B**, Forecast error and predicted uncertainty as earlier resting visits are added. **C**, Gain from personal history relative to no history or matched-other history. **D**, Forecast error with the personal reference compared with population or matched-other references. **E**, Agreement between predicted and observed response differences in participants with similar day-0 compositions, using personal, population or shuffled references. **F**, Observed change grouped by predicted change, with personal or population references. **G**, Correlation between predicted and observed change for each cell type, with personal or population references.

Forecasts improved as the twin incorporated more of the individual’s observations (Supplementary Fig. 5). In 63 participants with at least four earlier resting visits, adding these visits one at a time reduced composition error by 27%, from 0.51 to 0.37, and the uncertainty reported by the twin decreased in parallel, remaining within 11% of the observed error at every step, so that the twin’s own uncertainty indicated how far each forecast could be trusted (Fig. 3B).

This advantage depended on whose observations were provided. With model weights and current inputs fixed, removing personal history lowered the cosine similarity of the forecast direction by 0.10 (*q* = 0.015), and replacing it with the history of another participant matched on age and current immune state lowered it by 0.05 (*q* = 0.017; Fig. 3C). The personal reference was similarly specific. Replacing it with the population reference, the average resting composition of training participants, increased composition error by 5%, with higher error in 51 of the 75 participants (*q* = 0.020), whereas replacing it with the reference of another participant matched on age, sex and current composition increased error by 25%, with higher error in 71 (*q <* 0.001; Fig. 3D). A reference borrowed from a participant who resembled the individual was thus five times as harmful as the population reference, indicating that it carries information specific to the individual that demographics and the current state do not capture (Supplementary Fig. 6).

Personal observations also separated participants who were alike at the time of forecasting. We identified the 278 pairs of participants with the most similar day-0 compositions, the closest 10% of all pairs among the 75 participants. Because the two members of each pair started from nearly the same state, differences between their forecasts arose mainly from their personal observations. These forecast differences matched the direction of the pair’s measured differences in change, with a mean cosine similarity of 0.22 using personal references, twice the 0.10 obtained with the population reference and more than three times the 0.06 obtained with personal references shuffled among participants (permutation *P <* 0.001; Fig. 3E). Forecasts and pair selection used one half of each day-0 sample’s cells and measured changes the other half, so that sampling noise shared between forecast and measurement could not create this match. Within individual cell types, the virtual cells generated by each twin also ranked participants by how they would change. When participants were divided into thirds by the predicted change of each cell type, those in the lowest third showed a measured decrease of 0.39 s.d. and those in the highest third an increase of 0.34 s.d.; this separation of 0.73 s.d. was twice the 0.36 s.d. obtained with population references (Fig. 3F). Predicted and measured changes were positively correlated in 20 of the 21 cell types, with a median correlation of 0.31 compared with 0.17 for population references, and correlated more strongly with personal than with population references in 18 (Fig. 3G). These analyses used the same split of cells as the paired comparison.

### Return to a personal immune reference

The dependence of these forecasts on personal observations raised the question of which individual features persist. Personal histories contained persistent individual features alongside transient responses to vaccination. Most cell subsets and plasma proteins retained their relative ordering between participants across two years (adjusted for age, sex, cytomegalovirus status and, for proteins, assay plate), whereas early responses, late responses and net changes over each season showed little cross-year correlation (Fig. 4A).

**Fig. 4.**
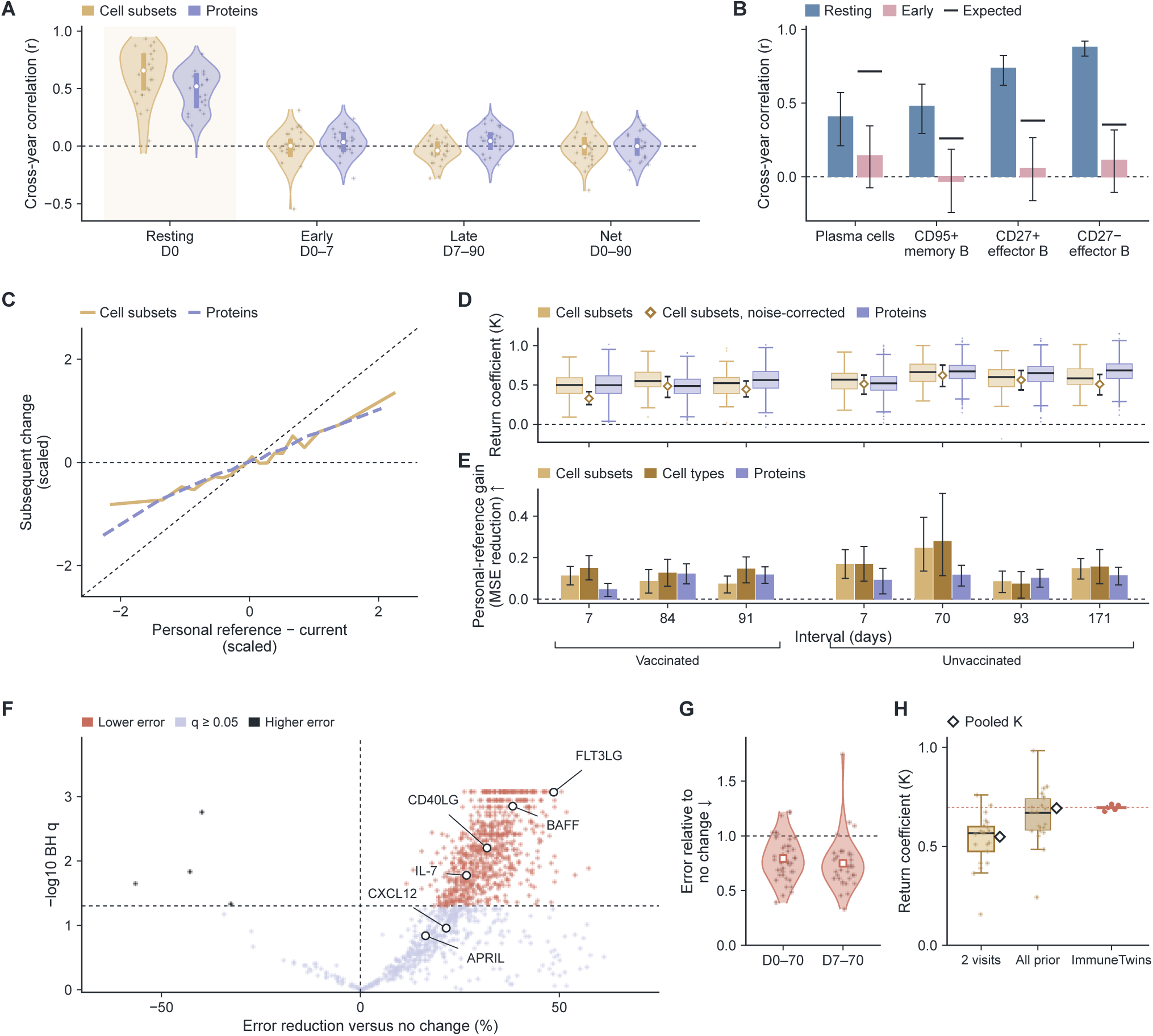
Persistent personal immune levels provide a reference for subsequent change. **A**, Cross-year correlations of resting levels and vaccine responses for cell subsets and proteins. **B**, Cross-year correlations of resting levels and early responses for B-cell subsets, with the correlation expected from measurement reliability. **C**, Subsequent change against deviation from the personal reference. **D**, Return coefficients across vaccinated and unvaccinated windows. **E**, Personal-reference gain across the same windows. **F**, The half-return rule compared with the no-change forecast for individual proteins. **G**, The half-return rule applied to flow-cytometry phenotypes in the independent NIH/CHI cohort. **H**, Return coefficients measured with two or all earlier resting visits as the reference and estimated by ImmuneTwins.

Against this stable background, plasma cells and CD27-positive effector B cells expanded during the first week in both seasons. Their resting levels remained correlated between years, whereas their early responses were much less correlated than expected for stable individual traits given the reliability of each year’s measurement (Fig. 4B). What persisted from one year to the next was therefore each individual’s level rather than the size of the response.

We then tested whether a participant’s personal level, estimated as the mean of the first-year resting visits, could serve as a reference for predicting subsequent change. In the second-year forecast window, deviations from this personal reference were followed by change towards it ^46^, with roughly half of the deviation recovered by day 90 for both cell subsets and proteins (Fig. 4C). This return occurred in all seven vaccinated and unvaccinated windows examined, spanning one week to nearly six months (Fig. 4D). For cell subsets, it persisted after correction for cell-sampling noise and was weakest in the first week after vaccination (Supplementary Fig. 7). In every window and feature set, forecasts based on the personal reference were more accurate than those based on the population reference (*q <* 0.05 for all 21 comparisons), distinguishing this return from regression towards the population mean (Fig. 4E).

ImmuneTwins forecasts with this return. Its personal-reference layer moves each forecast composition from the current state towards the individual’s reference by a return coefficient, which is estimated from training participants in each cross-validation fold. When the reference was defined as in the twin, from all earlier resting visits, the coefficient estimated by the twin agreed with the coefficient measured directly from participants’ compositions, and both exceeded the coefficient obtained with references from the two first-year visits alone (Fig. 4H). This difference reflects the reliability of the reference: a reference built from fewer visits is noisier and attenuates the measured return. The twin therefore returns each individual towards their own reference at the rate present in the data.

The return applied by the twin also held for measurements outside the single-cell data on which the twin was built. We applied it to plasma proteins and to an independent cohort with the return coefficient fixed in advance at one half, the value measured for cell subsets with first-year references (Fig. 4C). For proteins in the forecast window of Fig. 3A, this half-return rule was more accurate than the no-change forecast for 701 of the 1,229 proteins and less accurate for 4 (*q <* 0.05), with significant gains for four of six prespecified immune signalling proteins, FLT3LG, CD40LG, IL-7 and BAFF (Fig. 4F). In the independent vaccine cohort of the NIH Center for Human Immunology (NIH/CHI) ^7^, in which we analysed 113 immune phenotypes measured by flow cytometry and the personal reference came from a pre-vaccination visit, the same rule had lower error than the no-change forecast for 36 of 43 participants forecast from day 0 and for 32 of 36 forecast from day 7 (Fig. 4G). The return on which the twin’s forecasts rest is therefore not specific to its training data: it extended to circulating proteins and to an independent cohort measured on a different platform (Supplementary Fig. 8).

### B-cell remodelling in supercentenarians

Across CIMA, Immunobiology and OneK1K, the patient encoder of the base model weighted B cells increasingly relative to their abundance in older participants, whereas the relative weight of CD8 T cells declined with age (Fig. 5A). Both trends were significant in each cohort and persisted after adjustment for lineage fraction (*q <* 0.001; Supplementary Fig. 9), suggesting that B-cell states carry age-related information beyond B-cell numbers. We therefore examined B-cell composition in exceptional longevity, focusing on the states represented within the markedly smaller compartment. In the Hashimoto cohort ^58^, B cells accounted for on average 3% of peripheral blood mononuclear cells in seven supercentenarians (SC), compared with 15% in five controls. Yet this smaller compartment showed stronger expression of the secretory programme (Fig. 5B–D).

**Fig. 5.**
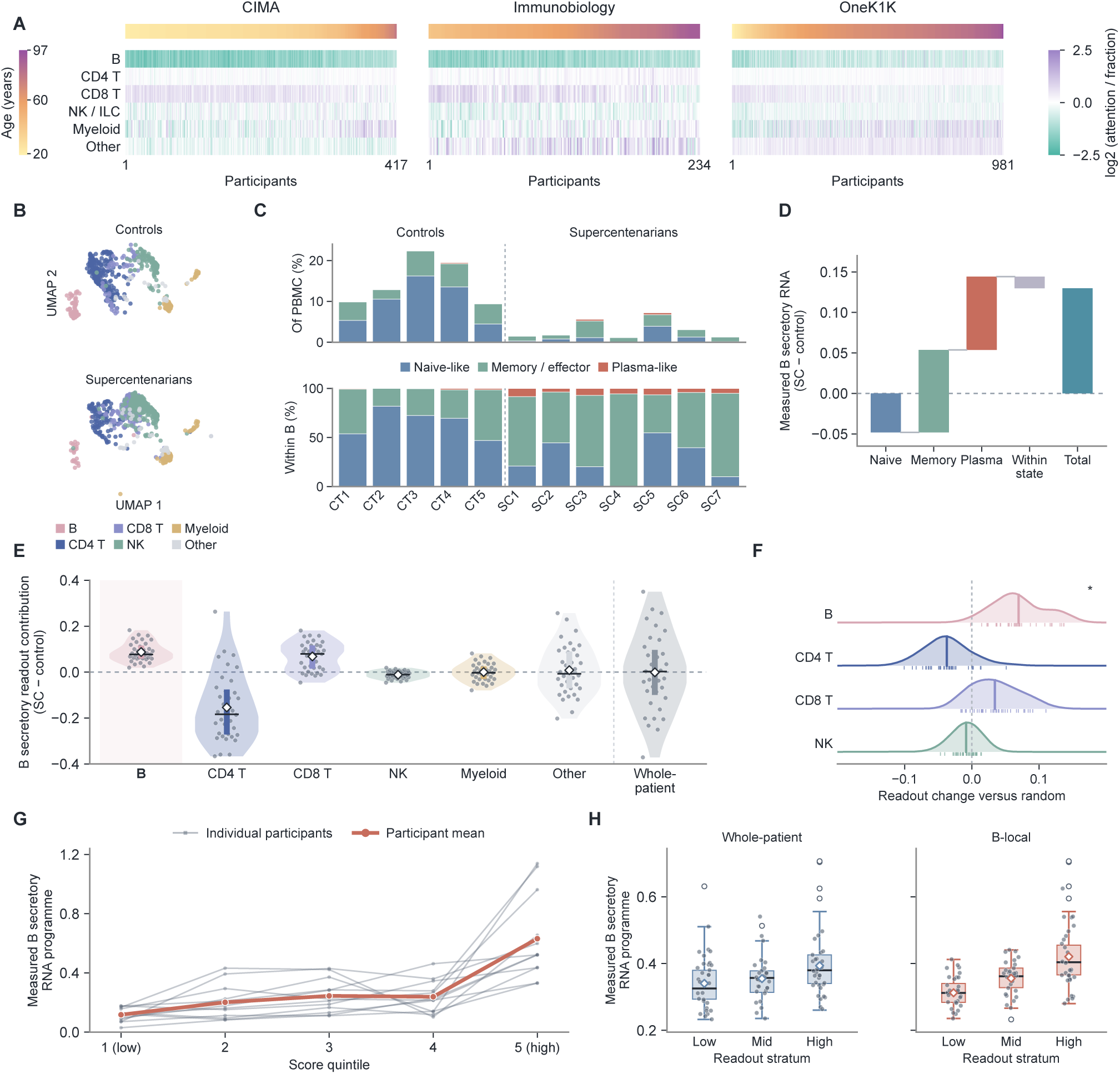
Cellular resolution of B-cell states in exceptional longevity. **A**, Lineage attention relative to cell fraction in three cohorts, with participants ordered by age. **B**, UMAP of cells from controls and supercentenarians. **C**, B-cell state composition in each control and supercentenarian. **D**, Decomposition of the group difference in the B secretory programme into state-allocation and within-state components. **E**, Contribution of each lineage to the B secretory readout. **F**, Readout change after replacing each lineage, relative to random replacement; the asterisk marks a significant difference. **G**, Measured B secretory programme across model-score quintiles within each participant. **H**, Measured B secretory programme across whole-patient and B-local readout strata in independent Immunobiology participants.

Decomposition of measured RNA attributed this difference to a shift in B-cell composition: naive-like states occupied a smaller fraction in supercentenarians, whereas memory/effector and plasma-like states were relatively enriched (Supplementary Fig. 10). This redistribution accounted for the higher mean expression of the secretory programme (Fig. 5C,D).

Counterfactual analysis with the twin identified which lineage produced this difference. For each of the 35 supercentenarian–control pairs, both participants were first reweighted to a common lineage composition, and the cells of each lineage were then exchanged between them in silico, so that the resulting change in the secretory readout could be attributed to that lineage. With the encoder and readout trained only in CIMA, B cells contributed more to the readout in supercentenarians than in controls (*q* = 0.015), whereas opposing contributions from other lineages offset this signal, so that the whole-patient readout barely differed between groups (Fig. 5E). Equal-count B-cell replacement produced a larger readout shift than lineage-matched random replacement (*q* = 0.041), indicating that the model captured differences in B-cell state rather than abundance (Fig. 5F).

Model readouts also corresponded to cellular expression: within each of the twelve participants, the highest-scoring B cells expressed a stronger secretory programme than the lowest-scoring cells (Fig. 5G). In 234 independent Immunobiology participants, B-local readouts likewise recovered individual variation in the programme more clearly than whole-patient readouts (Fig. 5H). Exceptional longevity thus showed a distinct B-cell configuration, with reduced representation accompanied by a shift towards memory and secretion-associated states within the remaining compartment. ImmuneTwins therefore resolved a change in B-cell state that the whole-patient read-out concealed: counterfactual analysis localized it to B cells, and the localized readouts agreed with measured expression within individuals and in an independent cohort.

### Immune trajectories during treatment

Unlike vaccination, myeloma induction therapy imposes a sustained perturbation, and no resting visits are available during treatment to estimate a personal reference; twins in this setting therefore relied on personal history alone. Among 20 patients receiving myeloma induction therapy ^3,59^, host immune trajectories differed even within the same tumour response category (Fig. 6A). Notably, CD4 T-cell fractions rebounded late in treatment, while B-cell fractions continued to fall and CD8 T-cell and natural killer-cell fractions decreased (Fig. 6F). Matched complete blood counts (CBC) from nineteen patients supported an increase in estimated CD4 T-cell counts during this period.

**Fig. 6.**
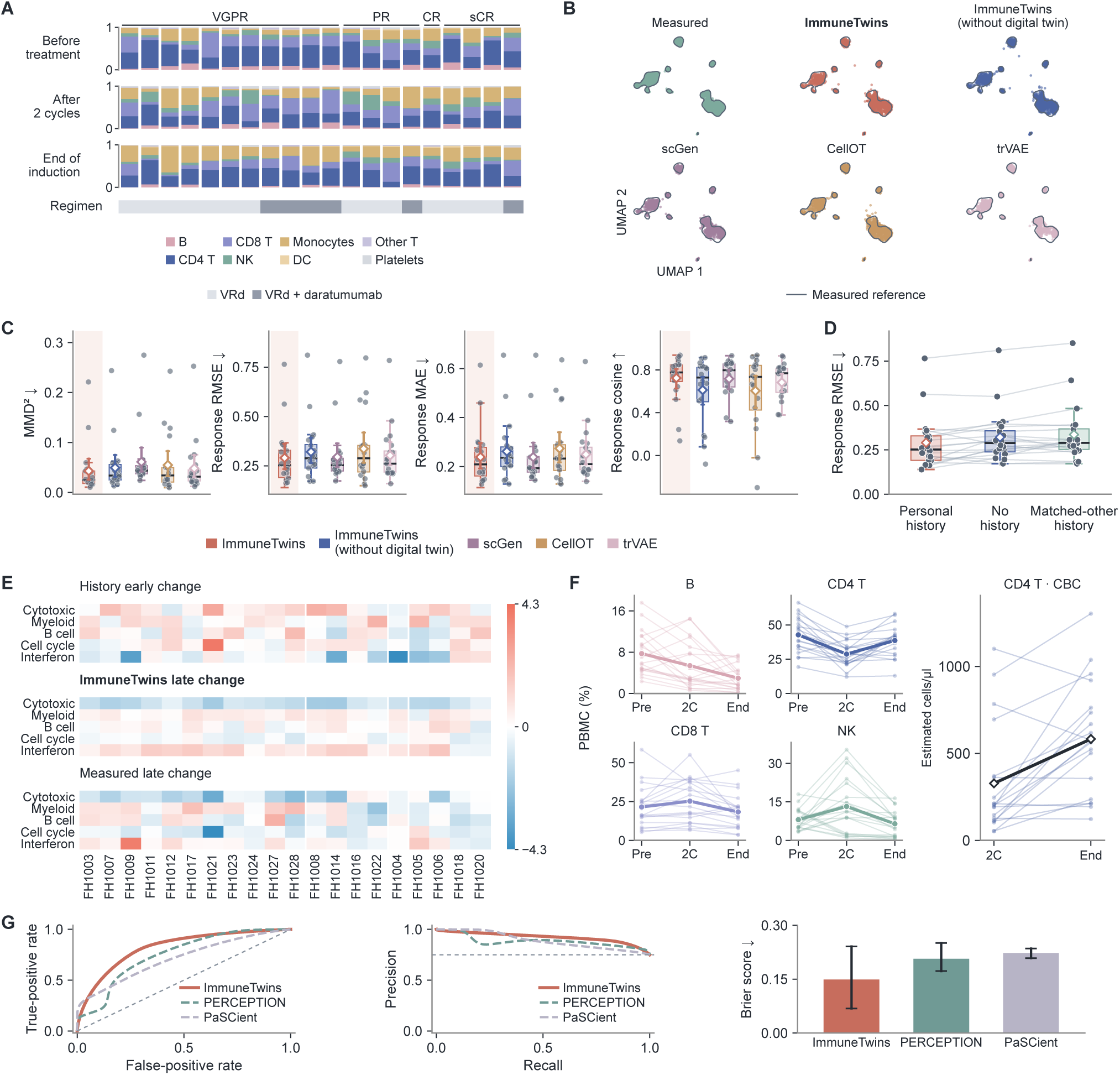
Personal immune trajectories during myeloma treatment. **A**, Lineage composition of each patient before treatment, after two cycles and at the end of induction, grouped by tumour response. **B**, Measured and predicted cell states of held-out patients in a common UMAP. **C**, Forecast accuracy of ImmuneTwins and other methods. **D**, Forecast error with personal history, no history or matched-other history. **E**, Early, predicted late and measured late programme changes in each patient. **F**, Lineage fractions and estimated CD4 T-cell counts during treatment. **G**, Prediction of six-month tumour response in the independent KYDAR cohort.

ImmuneTwins anticipated these later differences from each patient’s response over the first two treatment cycles, predicting the end-of-induction cellular distribution more accurately than the base model and published perturbation models ^10,49,57^ (Fig. 6B,C). This prediction depended on personal history: removing history or substituting that of another patient matched on regimen and current state increased prediction error (Fig. 6D). The forecasts captured individual differences in subsequent cytotoxic-programme decline and myeloid remodelling (Supplementary Fig. 11). Associations between predicted and measured changes persisted after adjustment for regimen and response grade (Fig. 6E), indicating that personal early responses provided information about later host immune changes beyond the tumour response category.

Baseline plasma-cell profiles also supported prediction of tumour response in the independent KYDAR cohort ^60^. ImmuneTwins predicted achievement of at least a partial response at six months, outperforming PERCEPTION ^61^ and PaSCient^38^ in both discrimination and response-probability prediction (Fig. 6G).

## Discussion

Here we present ImmuneTwins, a virtual-cell framework that builds a digital twin of each person’s immune system from single-cell profiles. Its patient representation, learned in one cohort, generalized zero-shot to populations of different ancestry and age, where it estimated immune age more accurately than existing methods. Twins that adapt a shared virtual-cell model to each person’s completed responses and personal reference forecast immune-cell changes after vaccination and during myeloma treatment more accurately than published methods.

These twins differ from earlier personalized predictions in how the individual enters the model. Studies of baseline immune states that predict vaccine responses ^7–9^ and virtual-cell models of perturbation responses ^5,10,49,50^ describe a person by a single measurement. Medical digital twins are personalized through organ physiology ^51^ or recorded history ^52^, and separating stable personal baselines from individual change has been proposed as a next step for longitudinal models ^53^. ImmuneTwins implements this separation in a virtual-cell model: personal history conveys how a person has responded, the personal reference conveys the state to which they return, and both enter a shared model as observed inputs. This changes how a virtual cell generalizes. Virtual cells still struggle in contexts not seen during training and often fail to outperform simple baselines ^6,43^. Each person is such a context, and what defines it is not contained in any single sample. Supplied as an observed record, this information lets a shared model adapt to a new individual without retraining, so that a twin can be built from a few visits, refined at each new visit and queried counterfactually.

Because the twins operate on cells, they can also locate where individual differences arise. Similar overall phenotypes can rest on different cellular foundations: in silico exchange of cells between participants traced the stronger secretory programme of supercentenarians to B cells shifted towards memory and secretion-associated states, consistent with immune ageing as a dynamic process rather than a steady decline ^2,62,63^, and patients with the same tumour response followed different host immune trajectories. Tracing such differences to cell states turns broad descriptions of immune enhancement or decline into testable biological questions.

We found that each person’s immune system has a characteristic state of its own: it persists across years, the immune system returns to it after vaccination ^46^ even though responses differ from year to year, and a closely matched individual cannot stand in for it. This challenges how immune health is judged. Blood counts are read against population reference ranges and immune age against people of the same age ^23–25^, comparisons that discard what is specific to the person. We propose that each person’s own state should serve as the reference, and virtual-cell digital twins make this practical. Built from a few blood samples and updated at every routine draw, a twin specifies where a person’s immune system should return after vaccination; extended with longer records to infection and drug treatment, it would reveal a failure to recover before disease becomes apparent. The same view recasts immune ageing: the convergence of older adults towards shared immune levels ^23^ may reflect drifting personal states, a weakening return to them, or both, and twins trained on records spanning decades are the means to tell these apart. The question for each person would then be not how old their immune system looks, but whether it still returns to its own state.

## Methods

### Data sources and preprocessing

We learned patient representations from single-cell profiles in the Chinese Immune Multi-Omics Atlas (CIMA; 417 participants) ^31^. Age prediction was evaluated in the Immunobiology of Aging cohort (234 participants) ^2,55^ and OneK1K (981 participants) ^29^. Longitudinal data included Sound Life samples collected on days 0, 7 and 90 around two annual influenza vaccinations and in an unvaccinated series, together with other visits outside these series ^2^, and samples from 20 patients receiving myeloma induction with bortezomib, lenalidomide and dexamethasone (VRd), with or without daratumumab, collected before treatment (Pre), after two cycles (2C) and at the end of induction (End) (GSE309595) ^3,59^. Tumour response in this cohort was graded as partial (PR), very good partial (VGPR), complete (CR) or stringent complete response (sCR). Additional analyses used Sound Life plasma proteins measured by proximity extension assay at the visits of the vaccinated and unvaccinated series, flow-cytometry phenotypes from the NIH/CHI influenza-vaccination cohort ^7^, seven supercentenarians and five controls from Hashimoto et al. ^58^ and 28 KYDAR participants with baseline plasma-cell profiles and clinical follow-up ^60^. Cells were grouped into B, CD4 T, CD8 T, natural killer (NK), myeloid and other lineages; the CIMA-based NK lineage included innate lymphoid cells (ILC), and treatment-cohort annotations separated monocytes, dendritic cells (DC), other T cells and platelets. Cohort sources and treatment visits are listed in Supplementary Tables 1 and 2.

Each visit was represented by an unordered cell set. The Universal Cell Embeddings model (UCE) was used to extract 1,280-dimensional features for each cell ^37^. Virtual cells were generated in a separate 16-dimensional RNA space: scVI coordinates for Sound Life ^64^ and linear coordinates fitted to log-normalized counts for the treatment cohorts. These coordinate systems were fitted using training participants and then applied to held-out participants. UCE and RNA cells were sampled independently within each visit and linked through the participant and sampling time. Transcriptional programme scores were calculated from predefined gene sets (Supplementary Table 3) by averaging normalized cell-level expression.

### Patient representation learning

The patient encoder projected UCE features to 128 dimensions and combined them by attention pooling ^65^. Attention weights were normalized across cells within each visit, and the weighted sum of projected features formed the patient representation. Pooling was invariant to cell order ^66^ and accommodated different cell counts ^67,68^. Projection and pooling parameters were learned jointly by predicting chronological age in CIMA, using a regression head and squared error on ages standardized within the fitting partition. UCE parameters remained fixed.

After model selection, the model was refitted on all CIMA participants for external application. The age head was retained for external age prediction. The cell projection and pooling layers were kept fixed in the virtual-cell model.

### Virtual-cell model

The base model of every twin was a virtual-cell model, consisting of the patient encoder and a response decoder shared by all individuals, that generated an individual’s future cells from their current cells. For each cell **z***_c_* of the current visit, the response decoder generated a virtual cell **y**^*_c_* at the target visit, conditioned on the current visit and response phase, and each virtual cell carried the cell-type label of the current cell from which it was generated. Cells were sampled independently at current and target visits. Learned query vectors (slots) attended separately to projected UCE features and RNA coordinates, followed by self-attention among the slots ^69^. Their pooled output, the patient representation and an encoding of the RNA mean, standard deviations and covariance formed a response context.

The response decoder first predicted mean and covariance changes within fixed response bases obtained by singular value decomposition of measured changes in training participants. Mean changes were expressed as a positive magnitude and a normalized direction in the mean-response basis; covariance changes used a separate basis of symmetric matrices. For current RNA coordinates **Z** *∈* R*^d×n^*, with cells as columns, mean ***µ*** and predicted mean change ***δ***, the affine prediction of the virtual cells was

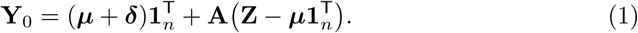

Here *d* = 16, *n* is the number of cells and **1***_n_* is the all-ones vector. The transform **A** was constructed from the current covariance and the predicted covariance change, with relative covariance eigenvalues constrained to training-derived bounds. Response coefficients were predicted from the response context.

The decoder then refined the shape of the virtual-cell distribution with a set network that updated whitened virtual-cell coordinates using local cell attention, pooled cell context and conditioning slots. After refinement, the coordinates were recentred and whitened again before restoring the affine prediction’s mean and covariance:

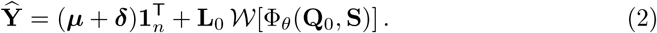

Here **Q**_0_ = *W*(**Y**_0_), **S** contains the conditioning slots, and Φ*_θ_* is the refinement network. The transformation *W* centres a cell set and whitens its unbiased covariance; **L**_0_ is the lower Cholesky factor of the empirical covariance of **Y**_0_. The final transformation preserves the affine mean and covariance while allowing changes in distribution shape.

### Personalizing the virtual-cell model

A digital twin personalized this virtual-cell model with two components: a history adapter that used the responses the individual had already completed (personal history) and a personal-reference layer that used the individual’s usual cell-type composition, estimated from earlier resting visits (personal reference). The history adapter used transitions completed by the current visit. Each transition was encoded from its initial and final mean RNA states, their difference, response phase, duration and order. A transformer encoded the transition sequence ^70^. A query derived from the unadapted current-state response context attended to these encodings to obtain the history embedding **h**. This summary controlled low-rank residual maps at adaptation site *j*:

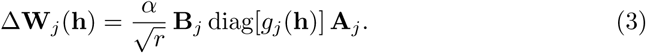

The shared matrices **A***_j_* and **B***_j_* defined a residual map through an *r*-dimensional bottleneck; *g_j_*was an affine projection followed by tanh, and *α* set the adaptation scale. Residual maps and additive prompts derived from history modified the slots and response context. Residual maps also adjusted the mean-direction, magnitude and covariance heads. The adapted slots were used for shape refinement. These history-dependent components were fitted with the base model fixed. At inference, personal history determined the adaptation without parameter updates; without history, the adapter left the virtual cells of the base model unchanged.

The personal-reference layer acted on the history-adapted prediction **Ŷ**. Each virtual cell **ŷ**_*c*_ was first rescaled as **ỹ***_c_* = **z***_c_* + *γ*(**ŷ***_c_* − **z***_c_*), where **z***_c_* is the current cell from which it was generated; the response scale *γ* thus changed the magnitude but not the direction of the predicted response. The layer then forecast the cell-type composition of the virtual cells, represented by centred log-ratio (CLR) transformed proportions of 21 cell types. For participant *i* with current composition **x***_i_*, the personal reference **x̅_*i*_** was the mean composition of the *n_i_* earlier resting visits, that is, visits other than those 7 days after vaccination. This reference was shrunk towards the population reference **x̅**^pop^, the mean resting composition of training participants, and the forecast moved from the current composition towards the shrunken reference **r̃***_i_*:

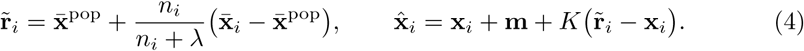

Here *K* is the return coefficient, **m** is the mean composition change of training participants over the forecast window and *λ* sets the strength of shrinkage; without earlier resting visits, the reference equals the population reference. The forecast variance, 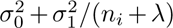, decreases as resting visits accumulate. Finally, the virtual cells were resampled by their cell-type labels to follow the composition softmax(**x̂**_*i*_). A twin was therefore personalized only through the individual’s own observations, with all parameters shared across individuals, and was updated by adding each new visit to its personal history or reference.

### Model training and evaluation

After representation learning, the affine response model, shape refinement and history adaptation were fitted sequentially, holding previously fitted components fixed. Forecasting losses compared virtual-cell and observed distributions and mean responses, with additional terms for response direction, magnitude and between-participant response geometry. History learning compared own-history response errors with errors obtained using shuffled histories and the fixed base model. Losses were averaged across seasons within each participant, then across participants.

Data were split by participant, keeping all visits and vaccination seasons from each participant in the same fold. Age prediction used five CIMA folds, giving out-of-fold (OOF) predictions within CIMA; external predictions were zero-shot, applying the model refitted on all CIMA participants without target-cohort calibration. Sound Life forecasting used five outer folds, with training duration selected within the outer-training participants before refitting. During inner selection of history-adaptation duration, the outer-fold RNA coordinates and base model were kept fixed. The personal-reference layer was then fitted in outer-training participants: *K* and **m** by least squares on measured compositions, *λ* and the variance terms by inner cross-validation, and *γ* from out-of-sample predictions of neural components refitted within inner folds.

The main vaccination task predicted second-year day-90 states from the second-year baseline, using first-year responses over days 0–7 and 7–90 as personal history and the resting visits before the baseline as the personal reference. Return to the personal reference was assessed in seven vaccinated and unvaccinated windows (Supplementary Table 4). For treatment forecasting, profiles after two cycles and the preceding treatment response were used to predict induction-end states without the personal-reference layer. Response parameters were fitted separately for each forecasting task. In KYDAR, a logistic regression head used representations of baseline plasma-cell profiles and forecast-derived features to predict whether a patient achieved at least a partial response at six months. Baseline-to-cycle-4 observations from training patients provided auxiliary supervision.

Age predictions were evaluated by mean absolute error (MAE), root mean squared error (RMSE), signed bias, Pearson correlation, the percentage within ten years, calibration slope and Lin’s concordance correlation coefficient (CCC) ^71^. Virtual-cell and observed target distributions were compared using squared maximum mean discrepancy (MMD^2^) ^72^. Mean-response errors and directions were assessed using RMSE, MAE and cosine similarity, and composition forecasts using the RMSE of CLR compositions, with equal participant weighting. History controls retained the fitted base model, history adapter and query cells while removing or substituting the history. Loss definitions, training procedures and evaluation details are given in Supplementary Notes 1 and 2, comparator implementations in Supplementary Table 5 and hyperparameters in Supplementary Table 6.

## Data availability

This study reanalysed existing datasets. The processed CIMA single-cell RNA-sequencing atlas and participant metadata are available from the CIMA Portal (https://db.cngb.org/trueblood/cima/resource) ^31^. Sound Life and Immunobiology of Aging single-cell profiles, plasma-protein measurements and clinical metadata are available from the Allen Institute’s Dynamics of Immune Health and Age resource (https://doi.org/10.57785/bn17-3h16) ^2^. OneK1K single-cell data are available from CZ CELLxGENE Discover (collection https://cellxgene.cziscience.com/collections/dde06e0f-ab3b-46be-96a2-a8082383c4a1)^29^. Supercentenarian single-cell count matrices and sample annotations are available from RIKEN SC2018 (http://gerg.gsc.riken.jp/SC2018/) ^58^. NIH/CHI influenza-vaccination flow-cytometry data are available from Zenodo (https://doi.org/10.5281/zenodo.10553454) ^7^.

Myeloma single-cell data are available from the Gene Expression Omnibus (GEO) under accessions GSE161195 (KYDAR) ^60^ and GSE309595 (longitudinal host immune profiling) ^3,59^; the matched blood counts, treatment records and tumour-response annotations for GSE309595 are available from the Allen Institute’s NDMM resource (https://apps.allenimmunology.org/aifi/insights/ndmm/downloads/clinical/). Numerical source data for Figs. 2–6 are provided with this paper.

## Code availability

The code used for data processing, model training and external validation is available at GitHub: https://github.com/Youzhuqinghuan/ImmuneTwins.

## Acknowledgements

This work was supported by the Prevention and Control of Emerging and Major Infectious Diseases–National Science and Technology Major Project (2025ZD01901502), the National Natural Science Foundation of China (62541160275), and the Natural Science Foundation of Yunnan (202504BW050004).

