## Supplementary Information for "Digital twins forecast individual immune trajectories"

#### Contents

**Supplementary Figure 1** **1**

**Supplementary Figure 2** **2**

**Supplementary Figure 3** **3**

**Supplementary Figure 4** **4**

**Supplementary Figure 5** **5**

**Supplementary Figure 6** **6**

**Supplementary Figure 7** **7**

**Supplementary Figure 8** **8**

**Supplementary Figure 9** **9**

**Supplementary Figure 10** **10**

**Supplementary Figure 11** **11**

**Supplementary Note 1** **12**

**Supplementary Note 2** **13**

**Supplementary Tables 1–6** **16**

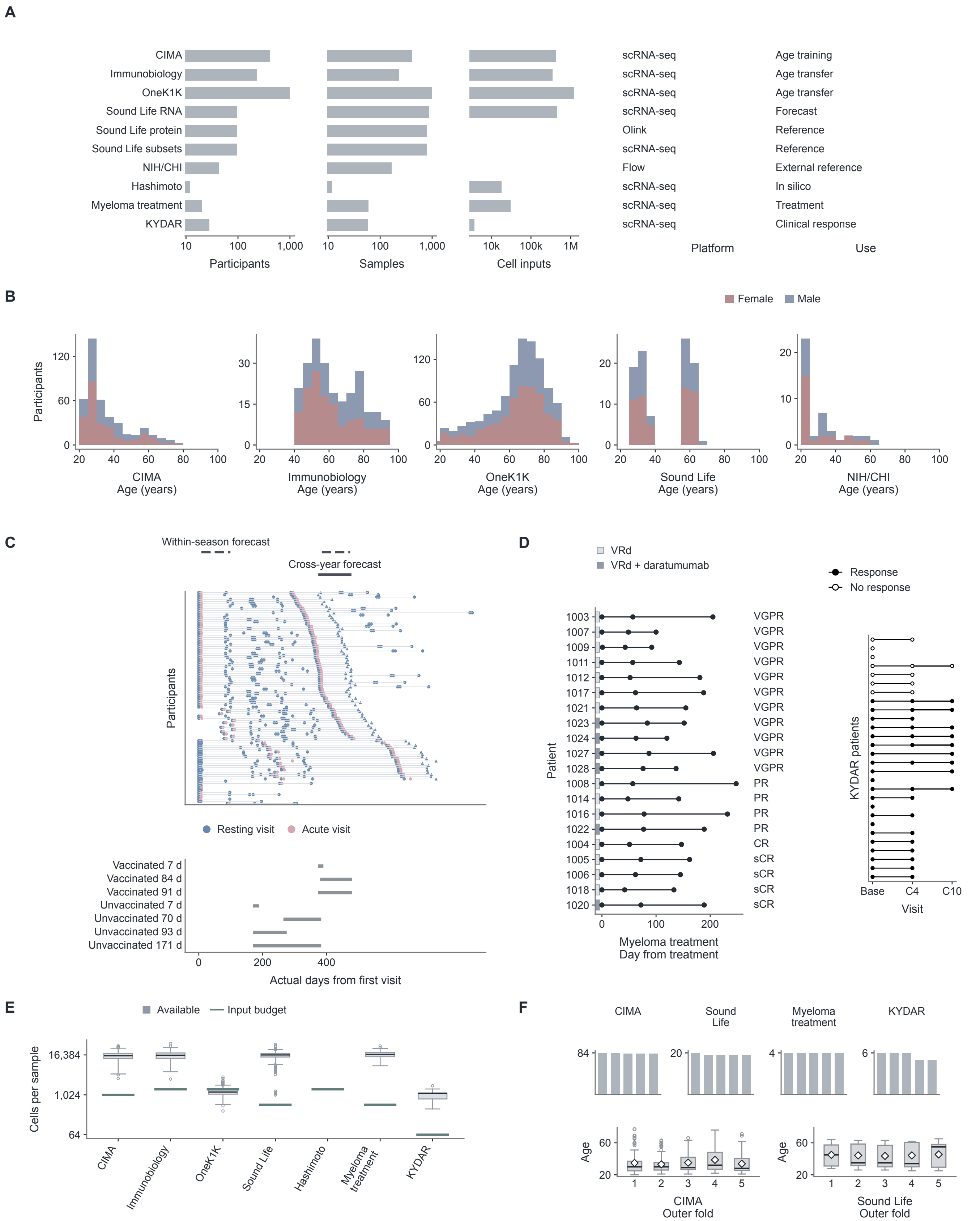


**Supplementary Figure 1 | Study cohorts, sampling schedules and model inputs.** **A,** Participants, samples and input cells for each cohort and measurement type, with platform and use in this study. **B,** Age distributions by recorded sex in CIMA (n = 417), Immunobiology (n = 234), OneK1K (n = 981), Sound Life (n = 96) and NIH/CHI (n = 43). **C,** Visit schedules of 95 Sound Life participants, with the cross-year and within-season forecast spans (top) and the seven vaccinated and unvaccinated windows (bottom). **D,** Visits, regimen and response grade of the 20 patients in the myeloma treatment cohort, and visits and response of the 28 KYDAR patients. **E,** Cells available per sample and cells used as model input in each cohort; per-sample totals were not available for Hashimoto. **F,** Participants in each outer cross-validation fold and age distributions within folds for CIMA and Sound Life. Boxes show medians and interquartile ranges, whiskers extend to 1.5 times the interquartile range and diamonds show means.

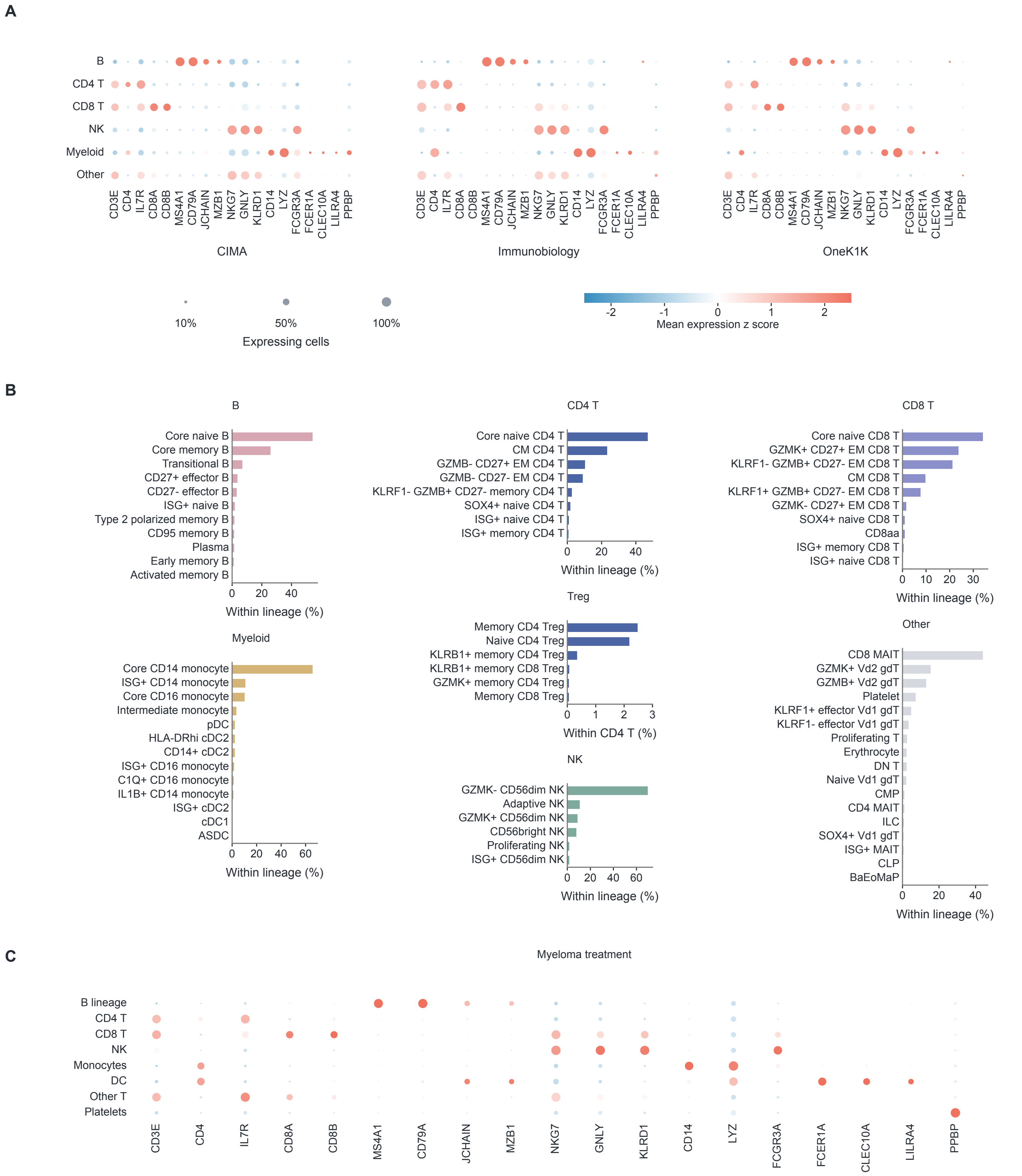


**Supplementary Figure 2 | Marker genes and lineage mapping used for cell annotation. A,** Expression of 19 marker genes across six major lineages in CIMA, Immunobiology and OneK1K. **B,** Assignment of 71 Sound Life cell subsets to major lineages, with the fraction of each subset within its lineage averaged across 95 participants; Treg subsets are shown as fractions of CD4 T cells. **C,** Expression of the same marker genes across eight lineages in the myeloma treatment cohort (20 patients, 60 visits). Dot size shows the fraction of expressing cells and colour the mean expression, z-scored across lineages.

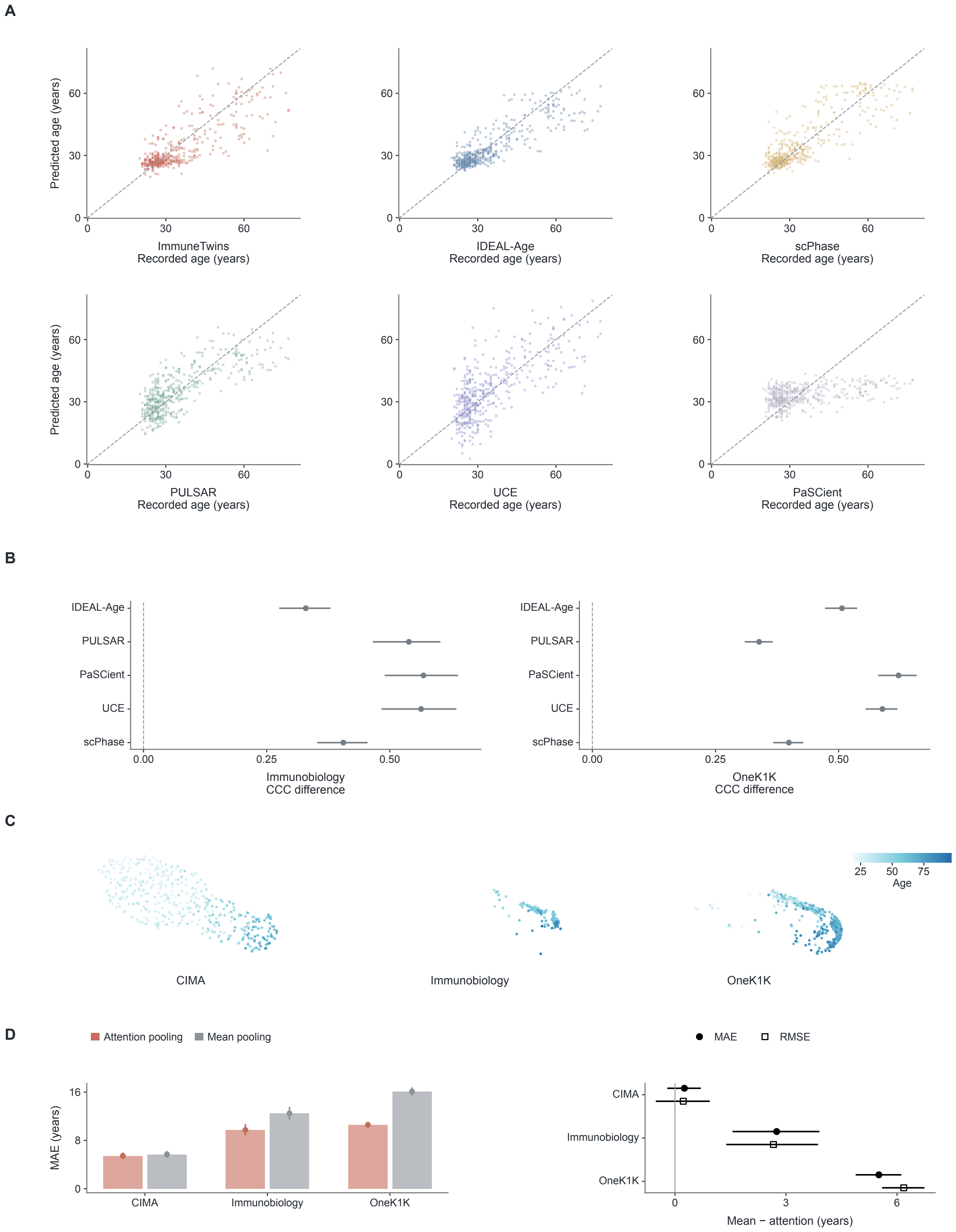


**Supplementary Figure 3 | Age prediction within CIMA and the contribution of attention pooling to transfer. A,** Out-of-fold predicted versus recorded age for six methods in CIMA (n = 417). **B,** Difference in concordance correlation coefficient between ImmuneTwins and each comparator in Immunobiology (n = 234) and OneK1K (n = 981). **C,** UMAP of patient representations from the three cohorts, coloured by recorded age. **D,** Age-prediction error with attention pooling or mean pooling, and the paired difference between them, in the three cohorts. Error bars show 95% confidence intervals from participant bootstrap resampling (10,000 resamples in B).

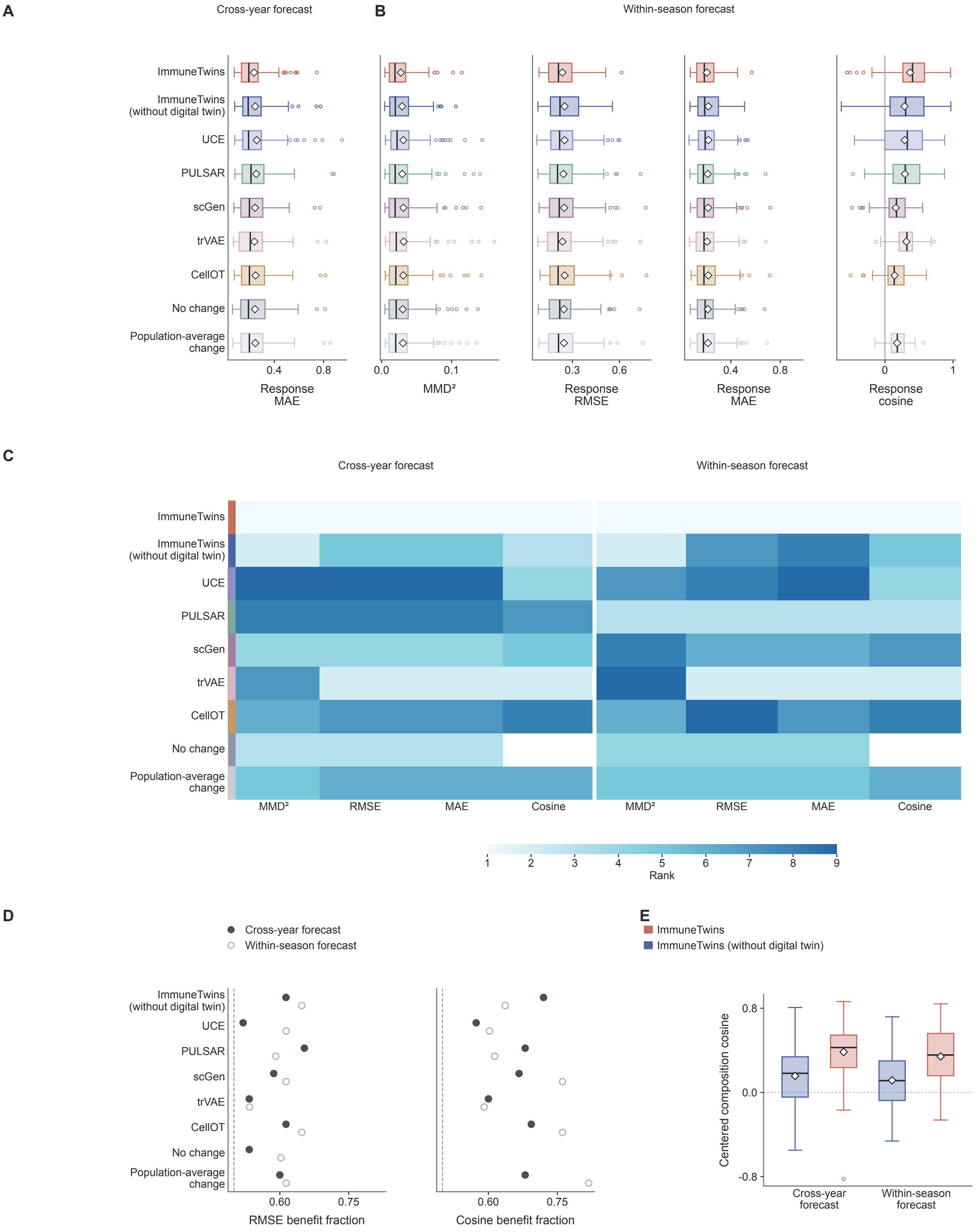


**Supplementary Figure 4 | ImmuneTwins ranks first on all RNA metrics for cross-year and within-season forecasts. A,** Response MAE of cross-year forecasts for nine methods ($n = 75$ participants). **B,** $MMD^2$, response RMSE, response MAE and response cosine of within-season forecasts from day 7 to day 90 ($n = 88$ participants, 163 participant-seasons). **C,** Rank of each method on each RNA metric in the two tasks; cosine is undefined for the no-change forecast. **D,** Fraction of participants for whom ImmuneTwins had lower error or higher cosine than each comparator. **E,** Agreement between predicted and observed composition changes after removing the mean change of training participants, with and without the digital twin. The digital twin increased this agreement by 0.23 in both tasks (95% confidence intervals 0.17–0.29 and 0.16–0.29; two-sided sign-flip tests, $q < 0.001$). Boxes show medians and interquartile ranges, whiskers extend to 1.5 times the interquartile range and diamonds show means.

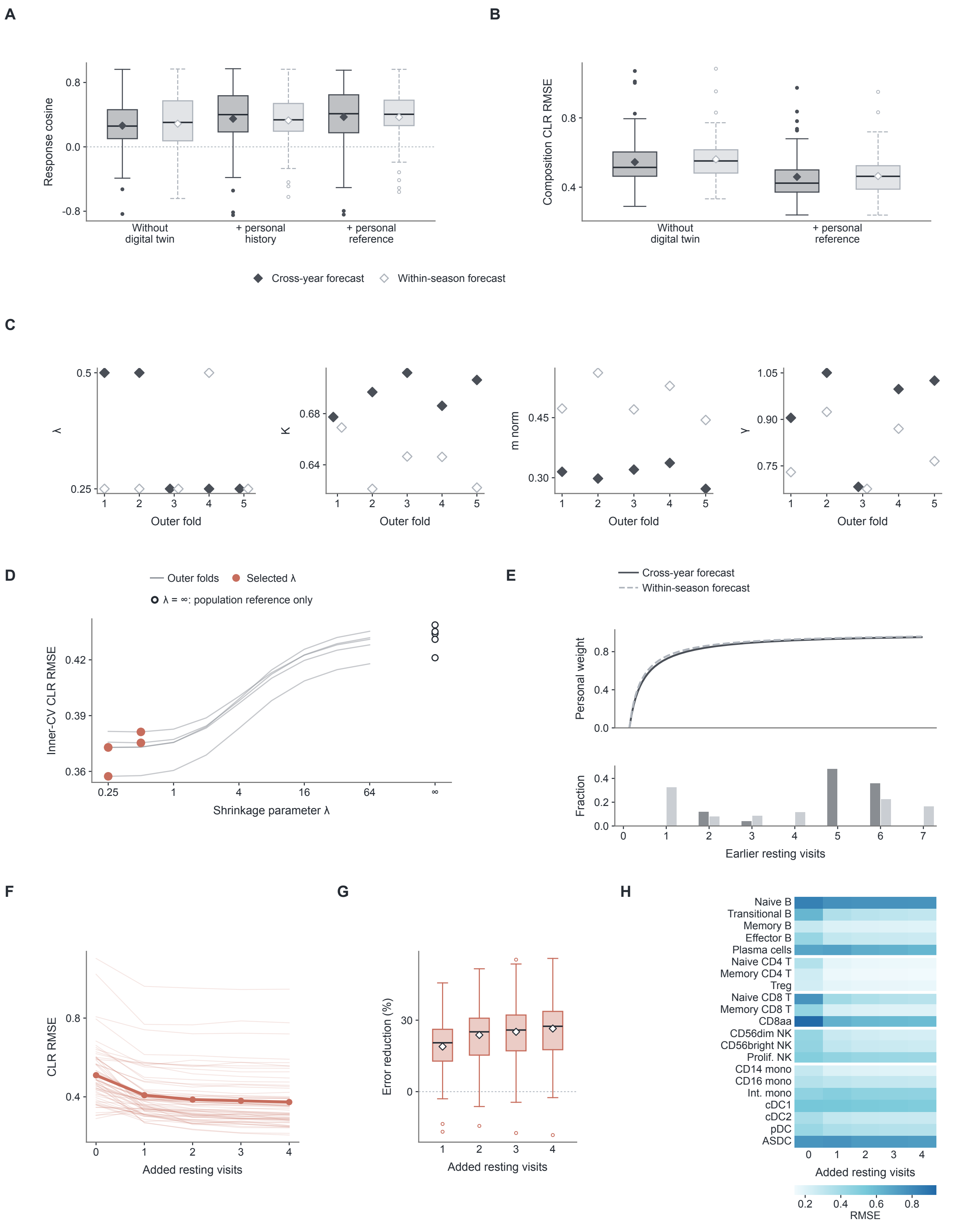


**Supplementary Figure 5 | Contributions of personal components, fitted parameters and additional visits to vaccine forecasts. A,B,** Response cosine (A) and composition error (B) as personal history and then the personal reference are added to ImmuneTwins (without digital twin), for cross-year ($n = 75$) and within-season ($n = 88$) forecasts. Personal history does not change composition forecasts and is therefore omitted from B; Fig. 3C instead removes history from the full twin. Benjamini–Hochberg-adjusted q values from two-sided paired sign-flip tests are $< 0.001$ and 0.097 (cross-year) and 0.052 and $< 0.001$ (within-season) for the history and reference steps in A, and $< 0.001$ in B. **C,** Shrinkage parameter $\lambda$, return coefficient K, mean displacement m and response scale $\gamma$ fitted in each outer fold. **D,** Selection of $\lambda$ by inner cross-validation for cross-year forecasts; $\lambda = \infty$ uses the population reference alone. **E,** Weight of the personal reference by number of earlier resting visits, with the distribution of visit numbers. **F,G,** Composition error (F) and its reduction (G) as earlier resting visits are added, in 63 participants with at least four such visits. **H,** Composition error of each cell type as visits are added. Boxes show medians and interquartile ranges, whiskers extend to 1.5 times the interquartile range and diamonds show means.

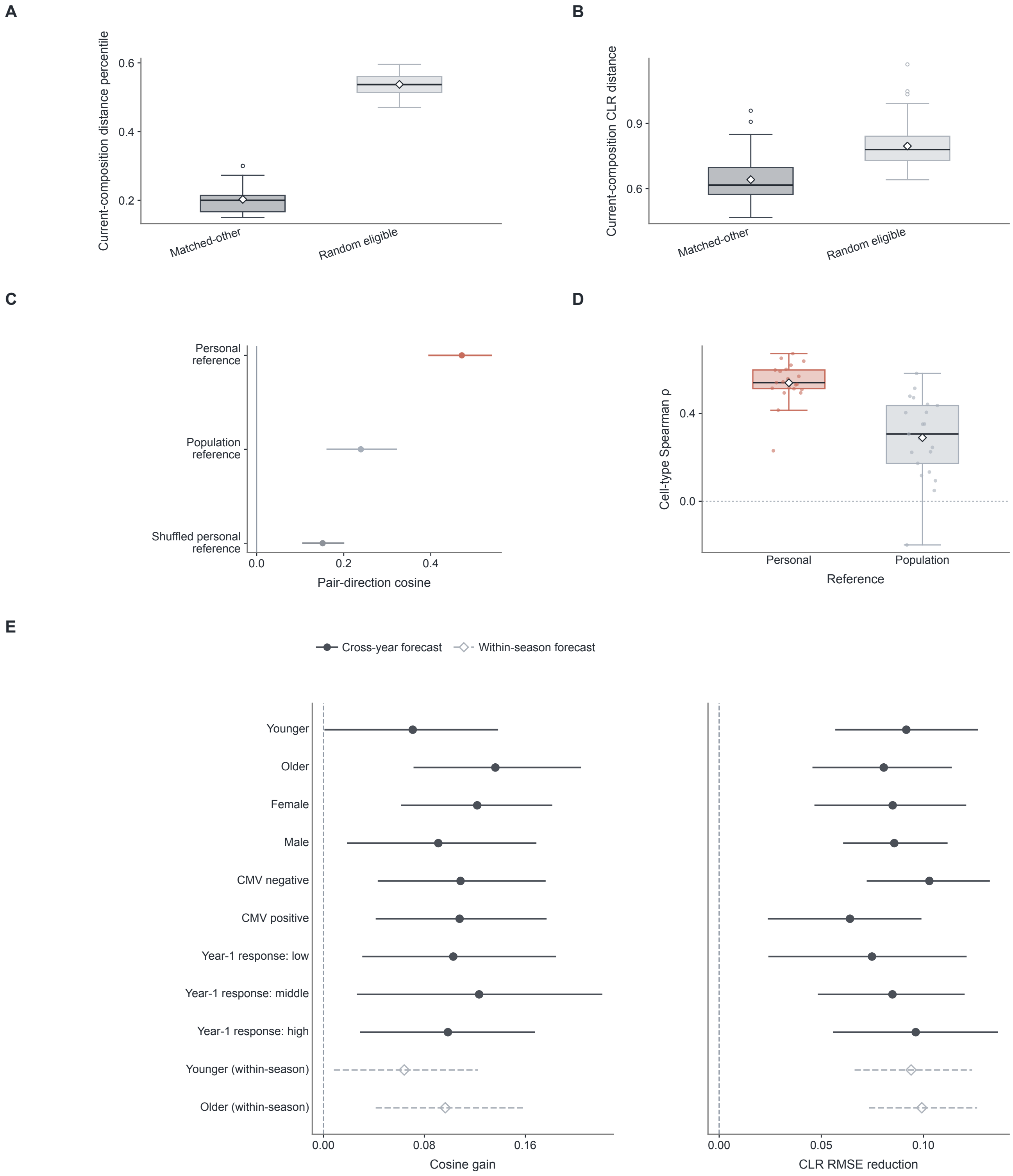


**Supplementary Figure 6 | The advantage of personal references persists with matched controls, without cell splitting and across subgroups. A,B,** Current-composition distance between each participant and the matched-other or a random eligible participant, as a percentile (A) and as CLR distance (B) ($n = 75$). Matched-other participants share age group and sex and have the closest current composition. **C,** Agreement between predicted and observed response differences in the 278 most similar participant pairs, using personal, population or shuffled references. **D,** Correlation between predicted and observed change for each of 21 cell types, with personal or population references. Unlike Fig. 3E,G, C and D use all cells of each sample rather than separate halves for forecast and measurement, which raises absolute values but leaves the order of references unchanged. **E,** Gain of ImmuneTwins over ImmuneTwins (without digital twin) in cosine similarity and composition error across participant subgroups of 25–45 participants. Error bars show 95% confidence intervals from 2,000 participant bootstrap resamples. Boxes show medians and interquartile ranges, whiskers extend to 1.5 times the interquartile range and diamonds show means.

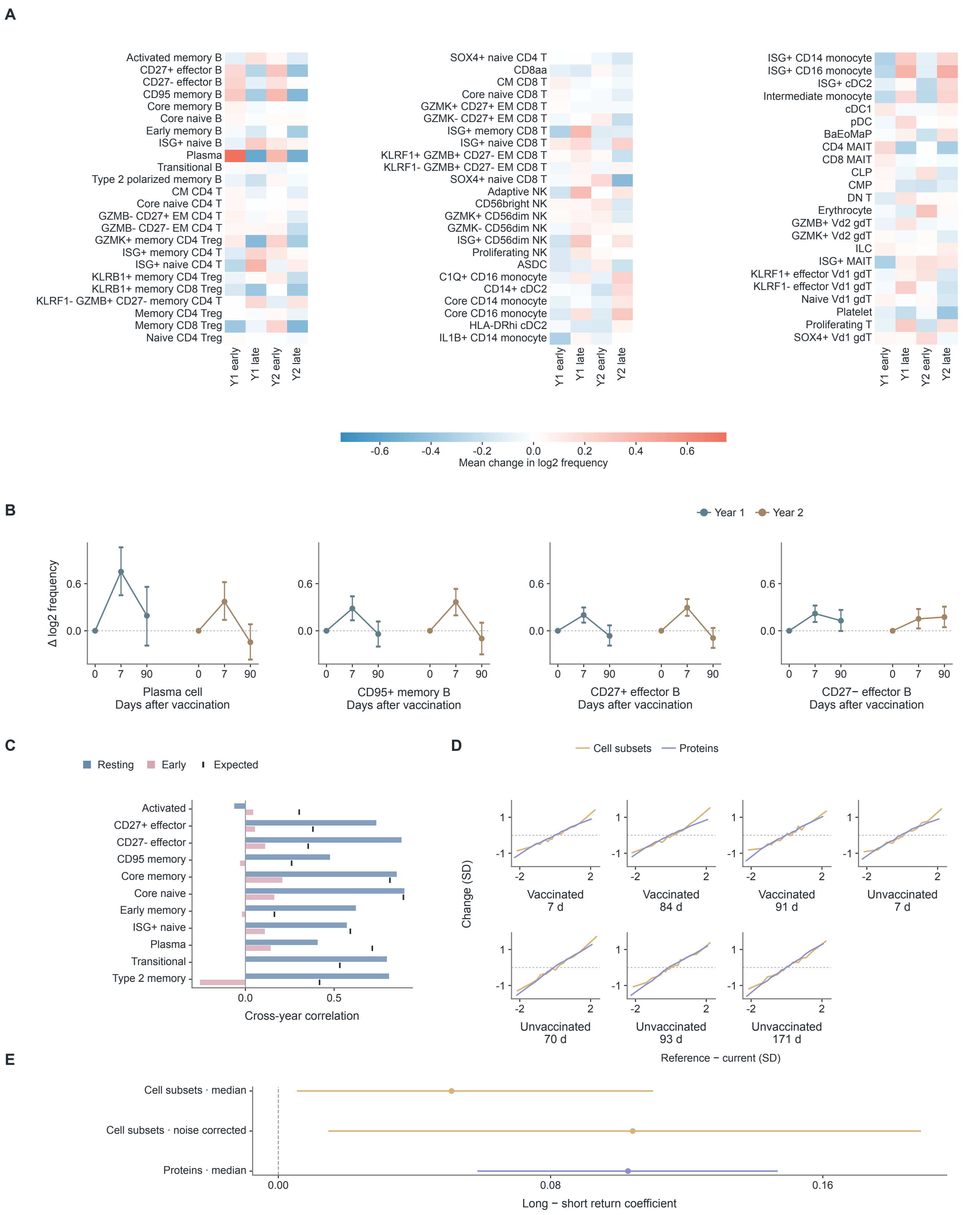


**Supplementary Figure 7 | Dynamics of fine cell subsets and dependence of the return on interval length. A,** Mean early (day 0–7) and late (day 7–90) changes of 71 cell subsets in each vaccination year (n = 75). **B,** Responses of plasma cells, CD95+ memory B cells and CD27+ and CD27− effector B cells in both years. **C,** Cross-year correlations of resting levels and early responses for all 11 B-cell subsets (n = 84), with the correlation expected from measurement reliability, extending Fig. 4B. **D,** Subsequent change against deviation from the personal reference for cell subsets and proteins in each of the seven windows (64–84 participants). **E,** Difference in return coefficient between the five longer windows and the two one-week windows; one-sided bootstrap P = 0.012 and 0.009 for cell subsets (median and noise-corrected estimates) and P < 0.001 for proteins. Error bars show 95% confidence intervals from participant bootstrap resampling.

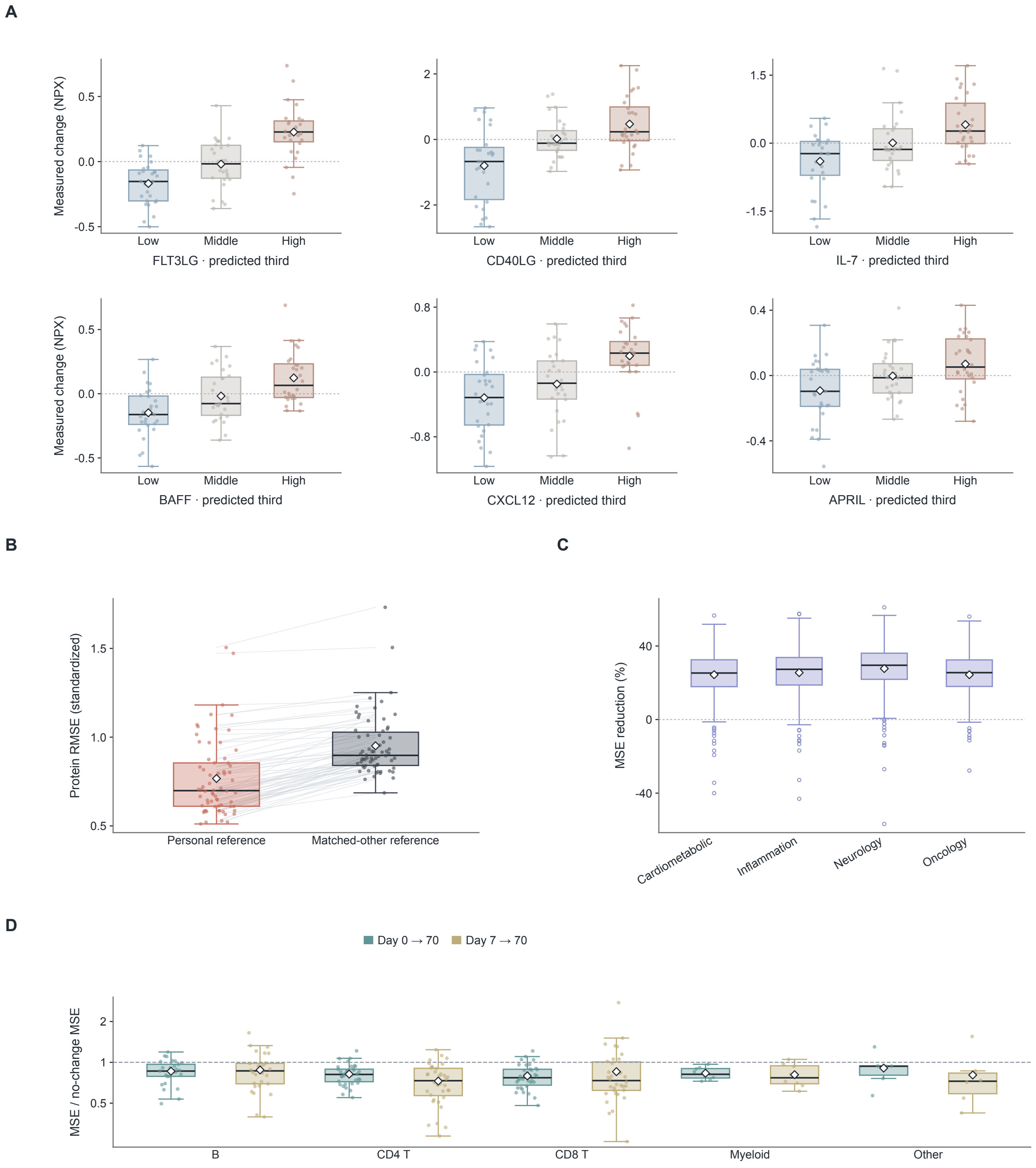


**Supplementary Figure 8 | The half-return rule extends to plasma proteins and to flow cytometry in an independent cohort. A,** Observed change in six prespecified immune signalling proteins, grouped by predicted change under the half-return rule (n = 82). **B,** Protein forecast error with the personal reference or the matched-other reference (n = 66). The personal reference reduced standardized MSE by 0.30 (95% confidence interval 0.27–0.34; two-sided sign-flip test, $P < 0.001$). **C,** Error reduction relative to the no-change forecast for 1,229 proteins, by Olink panel. **D,** Error of the half-return rule relative to the no-change forecast for 113 flow-cytometry phenotypes in the NIH/CHI cohort, by lineage, for forecasts from day 0 (up to 43 participants) and day 7 (up to 36 participants). Boxes show medians and interquartile ranges, whiskers extend to 1.5 times the interquartile range and diamonds show means.

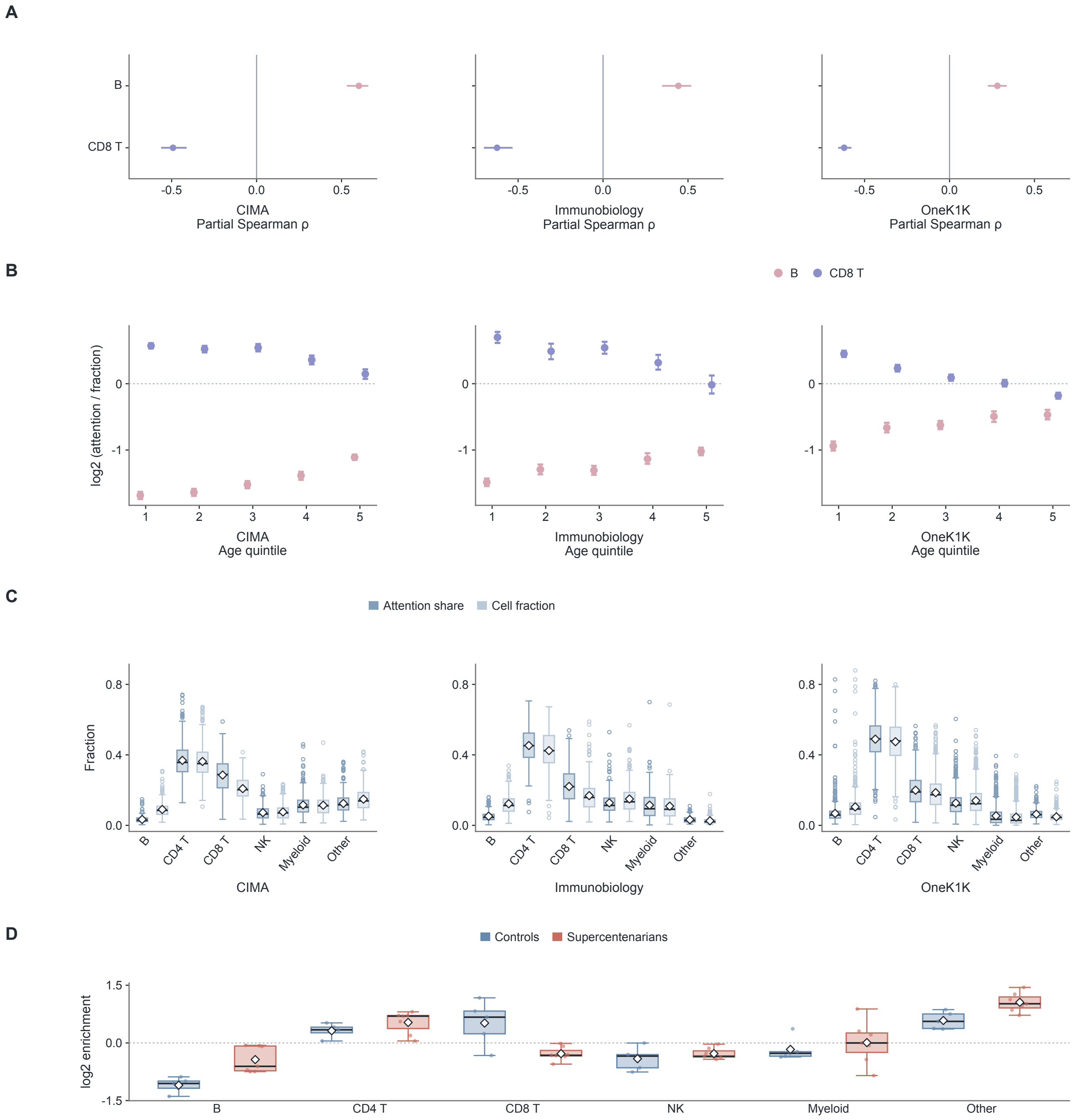


**Supplementary Figure 9 | Age-related attention to B and CD8 T cells persists after adjustment for their abundance.** **A,** Partial correlation between age and the attention share of B and CD8 T cells, adjusted for their cell fraction, in CIMA ($n = 417$), Immunobiology ($n = 234$) and OneK1K ($n = 981$). All six correlations had $q < 0.001$ (Benjamini–Hochberg correction across 18 cohort–lineage tests). **B,** Lineage attention relative to cell fraction for B and CD8 T cells across age quintiles. **C,** Attention share and cell fraction of the six lineages. **D,** Lineage attention relative to cell fraction in seven supercentenarians and five controls. Error bars show 95% confidence intervals from 2,000 participant bootstrap resamples. Boxes show medians and interquartile ranges, whiskers extend to 1.5 times the interquartile range and diamonds show means.

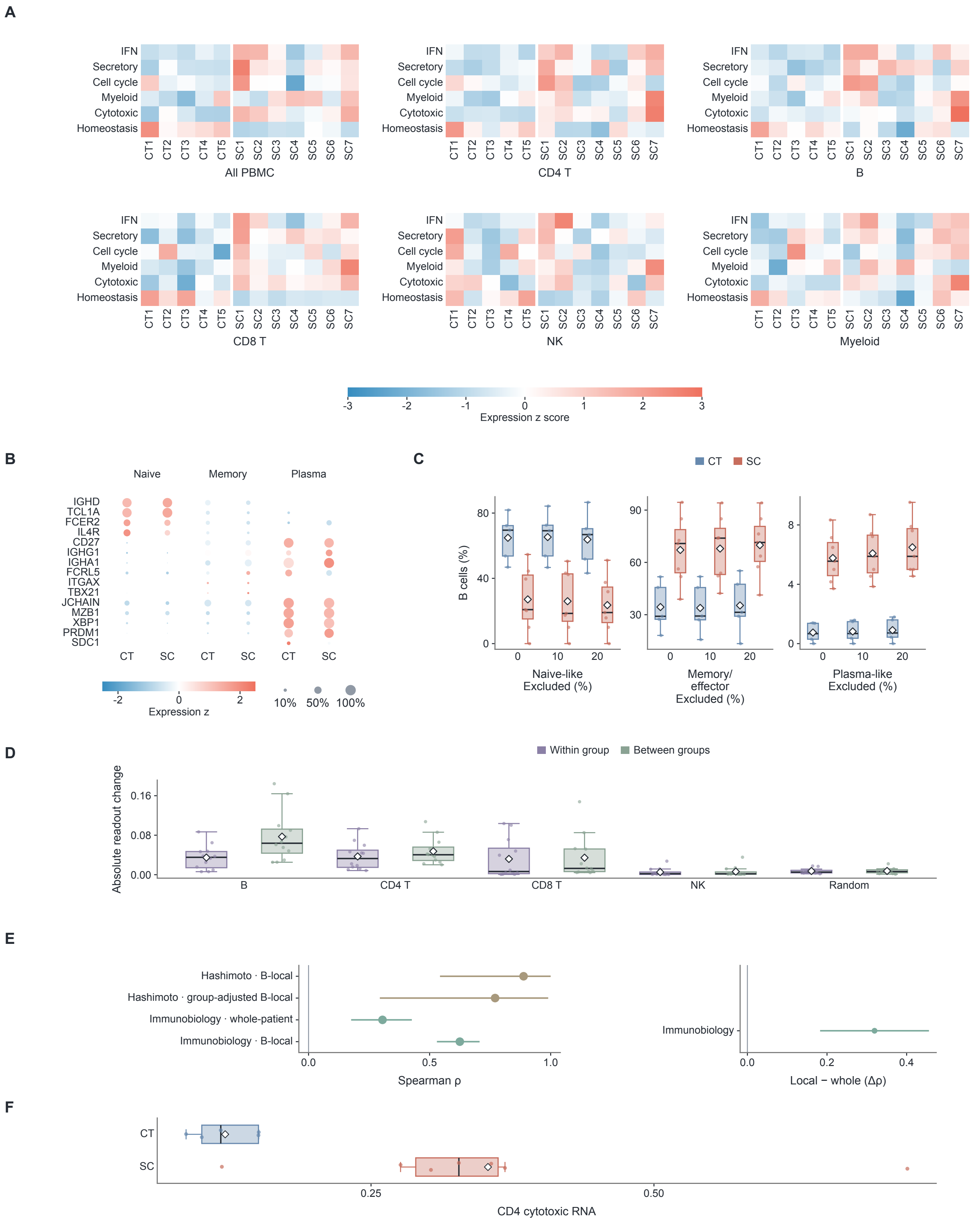


**Supplementary Figure 10 | Measured programmes, B-cell states and cell-replacement controls in supercentenarians.** **A,** Six RNA programmes in all PBMCs and in five lineages of seven supercentenarians (SC) and five controls (CT). **B,** Marker genes of naive-like, memory/effector and plasma-like B-cell states. **C,** B-cell state composition after excluding the 0%, 10% or 20% of B cells with the lowest annotation confidence. **D,** Change in the B secretory readout after replacing each lineage, or random cells, with cells from donors of the same or the other group. **E,** Correlation of B-local readouts (with and without adjustment for group) and whole-patient readouts with the measured B secretory programme in Hashimoto donors ($n = 12$) and Immunobiology participants ($n = 234$), and the B-local minus whole-patient difference. Error bars show 95% confidence intervals from 10,000 bootstrap resamples. **F,** Cytotoxic programme in CD4 T cells, reproducing the expansion of cytotoxic CD4 T cells reported in the original study (two-sided exact permutation test, $P = 0.010$; $q = 0.019$). Boxes show medians and interquartile ranges, whiskers extend to 1.5 times the interquartile range and diamonds show means.

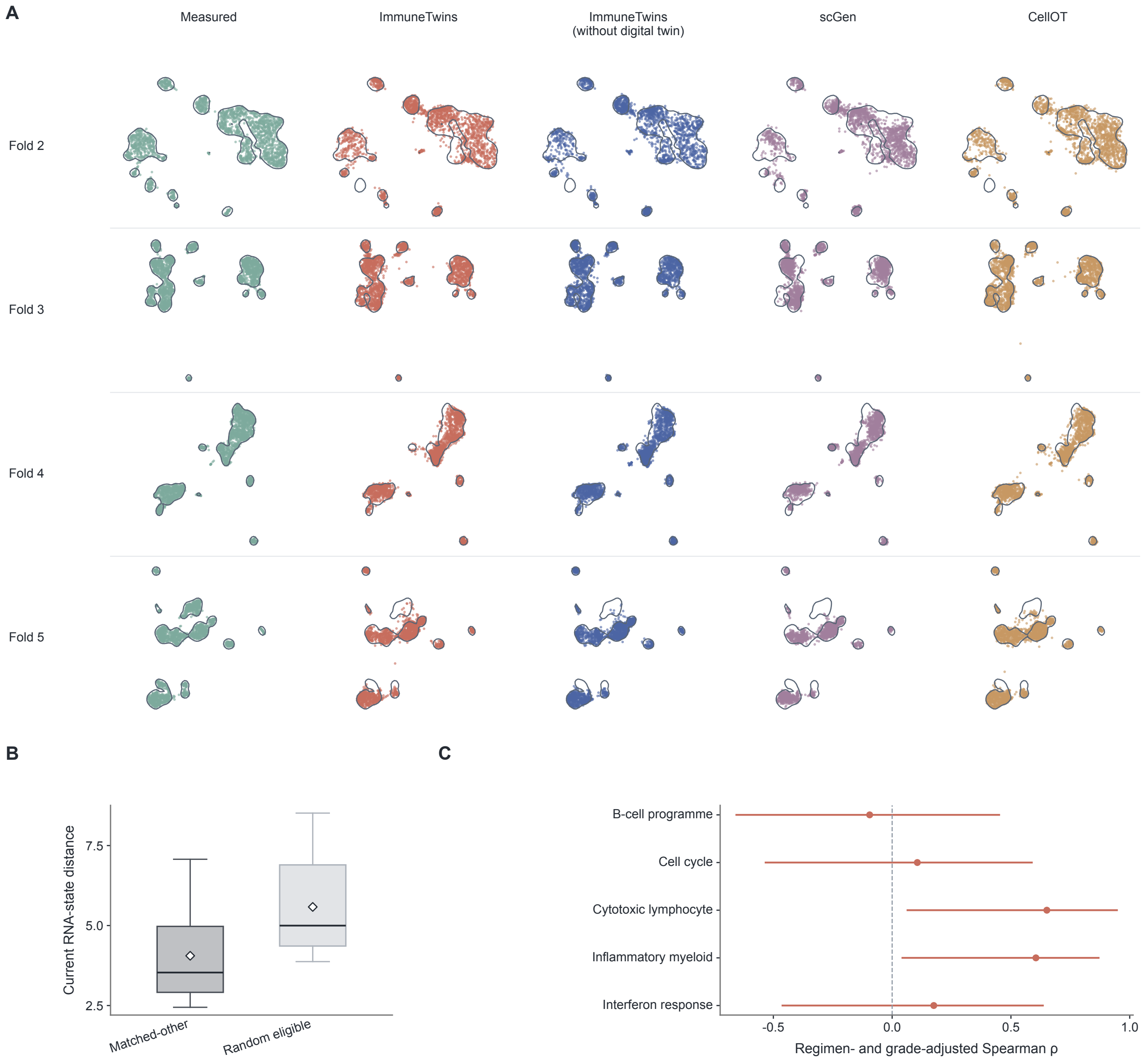


**Supplementary Figure 11 | Treatment forecasts in the remaining folds, matched histories and adjusted programme associations. A,** Measured and predicted end-of-induction cell states of the 16 held-out patients in folds 2–5, in a UMAP fitted to the measured cells of each fold; the remaining fold is shown in Fig. 6B. Outlines enclose 85% of the measured-cell density. **B,** Current RNA-state distance between each patient and the matched-other or a random eligible patient ($n = 20$); matched-other patients received the same regimen. **C,** Correlation between predicted and measured late programme changes, adjusted for regimen and response grade ($n = 20$), with 95% confidence intervals from 10,000 bootstrap resamples. Two-sided permutation tests with Benjamini–Hochberg correction gave $q = 0.032$ for the cytotoxic lymphocyte and inflammatory myeloid programmes and $q = 0.758$ for the other three. Boxes show medians and interquartile ranges, whiskers extend to 1.5 times the interquartile range and diamonds show means.

### Supplementary Note 1. Model construction and training

#### Response bases and affine prediction

RNA coordinates are scVI coordinates for Sound Life and linear coordinates fitted to log-normalized counts of training participants for the treatment cohorts. The mean- and covariance-response bases are the uncentred singular vectors of mean changes and of upper-triangular covariance changes in training transitions, each direction scaled by the root-mean-square of its training scores. The response context predicts the mean change $\boldsymbol{\delta}$, as a positive magnitude and a unit direction, and the covariance change $\Delta\mathbf{C}$. For current covariance $\mathbf{C}$ and regularizer $\epsilon$,

$$\mathbf{H} = (\mathbf{C} + \epsilon\mathbf{I})^{-1/2}(\mathbf{C} + \epsilon\mathbf{I} + \Delta\mathbf{C})(\mathbf{C} + \epsilon\mathbf{I})^{-1/2}, \qquad \mathbf{A} = (\mathbf{C} + \epsilon\mathbf{I})^{1/2}\,\mathrm{clip}(\mathbf{H})^{1/2}(\mathbf{C} + \epsilon\mathbf{I})^{-1/2}, \tag{S1}$$

where clip bounds the eigenvalues of $\mathbf{H}$ by the $q_{\mathrm{lo}}$ and $q_{\mathrm{hi}}$ quantiles of relative-covariance eigenvalues in training transitions, and $\mathbf{A}$ is the matrix in Methods equation (1).

#### Shape refinement and response scale

Each refinement layer combines local attention among virtual cells, a pooled summary of the virtual cells and attention to the slots, and updates whitened coordinates (Methods equation (2)). The response scale $\gamma$ is the participant-weighted least-squares coefficient of observed on predicted mean changes, fitted on inner out-of-fold predictions and clipped to the interval in Supplementary Table 6.

#### Personal history

Each completed transition is encoded from its source mean, target mean and their difference, with phase, duration and order embeddings. One Transformer layer encodes the transition sequence, and the unadapted response context queries it to give the history embedding $\mathbf{h}$. Low-rank residual maps (Methods equation (3)) and additive prompts derived from $\mathbf{h}$ modify the slots, the response context and the mean-direction, mean-magnitude and covariance heads; shape refinement receives the adapted slots. With no completed transition these terms are zero.

#### Personal reference

Resting visits are day-0, day-90, stand-alone and Immune Variation visits before the query visit. Counts $c_j$ of the $D$ cell types are transformed to centred log-ratio (CLR) coordinates,

$$x_j = \log(c_j + \varepsilon) - D^{-1}\sum_k \log(c_k + \varepsilon). \tag{S2}$$

The population reference is the participant-equal mean resting composition of training participants, and $\mathbf{m}$ is the participant-weighted mean composition change over the forecast window. For transition $u$, with current composition $\mathbf{x}_u$, shrunken reference $\tilde{\mathbf{r}}_u$ (Methods equation (4)) and observed target $\mathbf{y}_u$,

$$K = \frac{\sum_u w_u(\tilde{\mathbf{r}}_u - \mathbf{x}_u)^\top(\mathbf{y}_u - \mathbf{x}_u - \mathbf{m})}{\sum_u w_u\|\tilde{\mathbf{r}}_u - \mathbf{x}_u\|^2}, \tag{S3}$$

where $w_u$ gives each participant equal total weight. $\lambda$ is selected by inner out-of-fold CLR RMSE, and $\sigma_0^2$ and $\sigma_1^2$ are fitted to inner out-of-fold squared errors by non-negative least squares. Virtual cells are resampled systematically with weights equal to their type's forecast proportion, $\mathrm{softmax}(\hat{\mathbf{x}}_i)$, divided by the number of cells of that type and renormalized over the types present.

#### Losses

The distribution loss is the biased Gaussian MMD$^2$ (Supplementary Note 2). In standardized response coordinates, the mean loss is the squared error of the mean change, the direction loss is one minus the cosine similarity of predicted and observed changes, and the magnitude loss is the smooth-$L_1$ difference between $\log(1 + \|\boldsymbol{\delta}\|)$ and $\log(1 + \|\boldsymbol{\delta}_{\mathrm{obs}}\|)$. The geometry loss centres predicted and observed responses across the participants of each update, scales them by the root-mean-square observed response, and combines smooth-$L_1$ differences of the centred responses (weight 0.75) and of their Gram matrices divided by the RNA dimension (weight 0.25). Stage weights are:

| Stage | Distribution | Mean | Direction | Magnitude | Geometry |
|---|---|---|---|---|---|
| Affine response | 0.25 | 1 | 0.25 | 0.25 | 0.50 |
| Shape refinement | 1 | 1 | 0.25 | 0.25 | 0.50 |
| Personal history | 1 | 2 | 0.50 | 0.50 | 0.75 |

History training adds two losses comparing the response error with the participant's own history, $e_{\text{own}}$, with that obtained with a shuffled history, $e_{\text{shuffled}}$, and with that of the fixed base model, $e_{\text{base}}$:

$$\mathcal{L}_{\text{history}} = \text{softplus}[(e_{\text{own}} - e_{\text{shuffled}})/\tau_1], \qquad \mathcal{L}_{\text{anchor}} = \text{softplus}[(e_{\text{own}} - e_{\text{base}})/\tau_2], \tag{S4}$$

with weights 0.5 and 0.25. Response losses are averaged over transitions within each participant and then over participants.

##### Training and model selection

All visits of a participant belong to one outer fold, and all coordinates, scales and parameters are fitted on outer-training participants. The age model minimizes the squared error of standardized age, with checkpoints selected by inner validation error; external predictions use a model refitted on all CIMA participants for the median selected epoch, without recalibration. Components are trained in sequence with earlier components fixed: the affine response model is selected by validation response RMSE, shape refinement by validation $\text{MMD}^2$, and history adaptation by the rounded median of inner-fold best epochs (within season) or the minimum of the participant-weighted validation curve averaged across inner folds (cross year), before refitting on all outer-training participants.

For treatment, the pretreatment-to-two-cycle transition is the history, a shared recovery basis describes the second phase and no personal reference is used; the history learning rate and epoch minimize the patient-weighted validation response RMSE. For KYDAR, the response model is trained on baseline-to-cycle-4 transitions of training patients and forecasts from baseline without history. The logistic regression head uses the patient representation, current RNA mean, predicted mean change and radial-basis summaries of the current distribution and predicted change, with anchors and bandwidth fitted on training baseline cells; regularization and state-feature scaling maximize AUROC minus 0.2 times the Brier score in response-stratified inner folds.

#### Supplementary Note 2. Evaluation and analyses

##### Data selection and annotation

CIMA contributes one visit per participant, and external age analyses include participants with recorded exact ages. Labels are mapped in order to CD4 T (CD4 and regulatory T), CD8 T, B (B and plasma), NK (natural killer and innate lymphoid), myeloid (monocytes, dendritic cells and macrophages) and Other. Sound Life analyses use annotated cell types, 71 fine subsets and plasma proteins measured at both visits of each window, without imputation. NIH/CHI analyses use the 113 released flow-cytometry gates and require at least ten participants with complete measurements per forecast. Hashimoto analyses use primary samples, excluding technical repeats.

##### Forecast scoring

Within-season forecasts predict day 90 from day 7, and cross-year forecasts predict second-year day 90 from second-year day 0; only transitions completed before the query visit enter the history. RNA scores average 20 draws of 256 cells without replacement, then seasons within participants, then participants. Coordinates use outer-training centring and scaling, and all methods share one Gaussian bandwidth $b$ per outer fold, the median distance between outer-training cell pairs. With $k_b(\mathbf{u}, \mathbf{v}) = \exp[-\|\mathbf{u} - \mathbf{v}\|^2/(2b^2)]$ and cell sets $\{\mathbf{u}_c\}_{c=1}^{N}$ and $\{\mathbf{v}_c\}_{c=1}^{M}$,

$$\text{MMD}^2 = \frac{1}{N^2}\sum_{c,c'} k_b(\mathbf{u}_c, \mathbf{u}_{c'}) + \frac{1}{M^2}\sum_{c,c'} k_b(\mathbf{v}_c, \mathbf{v}_{c'}) - \frac{2}{NM}\sum_{c,c'} k_b(\mathbf{u}_c, \mathbf{v}_{c'}). \tag{S5}$$

For predicted and measured mean changes $\boldsymbol{\delta}$ and $\boldsymbol{\delta}_{\text{obs}}$ in $d$ dimensions, response RMSE, MAE and cosine similarity are

$$\sqrt{\|\boldsymbol{\delta} - \boldsymbol{\delta}_{\text{obs}}\|^2/d}, \qquad d^{-1}\|\boldsymbol{\delta} - \boldsymbol{\delta}_{\text{obs}}\|_1, \qquad \frac{\boldsymbol{\delta}^\top \boldsymbol{\delta}_{\text{obs}}}{\|\boldsymbol{\delta}\|\|\boldsymbol{\delta}_{\text{obs}}\|}. \tag{S6}$$

The no-change forecast is omitted from cosine comparisons. Composition RMSE is taken over CLR coordinates. Age predictions are evaluated by MAE, RMSE, mean error, Pearson correlation, fraction within ten years, slope of predicted on recorded age and

$$\mathrm{CCC} = \frac{2\,\mathrm{cov}(a,\widehat{a})}{\mathrm{var}(a) + \mathrm{var}(\widehat{a}) + (\bar{a} - \bar{\widehat{a}})^2}, \tag{S7}$$

with population moments. KYDAR is evaluated by AUROC, average precision and Brier score for at least a partial response; PERCEPTION Brier scores use nested patient-held-out Platt probabilities.

#### Substitution controls, updating and split halves

Matched-other histories come from the five held-out participants of the same fold with the closest current RNA states among those matched on age group, sex and cytomegalovirus status, relaxed stepwise to age group and sex, age group, and all participants. Matched-other references come from the five outer-training participants of the same season, sex and age group with the closest current CLR compositions. Errors are averaged over the five substitutes; query cells and parameters are fixed. Treatment histories are matched on regimen, then current RNA state. Updating draws 20 subsets of earlier resting visits at each history length from zero to four, with fitted parameters fixed. For participants with similar current states, half of the cells of each day-0 sample define current composition, pairing and forecasts, and the other half define measured change. The pairwise cosine of predicted and measured response differences is tested by permuting reference labels with pairs fixed, $P = (1 + \#\{s_{\mathrm{perm}} \geq s_{\mathrm{obs}}\})/(B+1)$ for $B$ permutations. Terciles and standardization are defined within each cell type.

#### Persistence and reference return

Cross-year correlations are adjusted for age, sex and cytomegalovirus status, and for proteins also for assay plate. Split-half reliability uses the Spearman–Brown correction, $2r_{\mathrm{half}}/(1 + r_{\mathrm{half}})$, and the correlation expected for a stable trait is $\sqrt{\max(0, \mathrm{Rel}_1)\max(0, \mathrm{Rel}_2)}$. Personal references in the return analyses average the first-year day-0 and day-90 visits. Return coefficients regress subsequent change on deviation from the reference, pooled across features by summing covariance and variance terms. With independent cell halves $A$ and $B$,

$$K_{\mathrm{noise}} = \frac{\sum_f \mathrm{cov}(y_f - x_{f,B},\, r_{f,A} - x_{f,A})}{\sum_f \mathrm{cov}(r_{f,A} - x_{f,A},\, r_{f,B} - x_{f,B})}. \tag{S8}$$

Personal and population references are compared by five-fold participant cross-fitting, with errors divided by the standard deviation of change in training participants. Protein and NIH/CHI forecasts use $\widehat{y} = x + 0.5(r - x)$. NIH/CHI gates are $\log_2$-transformed and forecast at day 70, with the day $-7$ visit as reference for day-0 forecasts and the mean of days $-7$ and 0 for day-7 forecasts; errors are scaled by pre-vaccination variation and divided by the no-change error.

#### Attention, cell replacement and programme readouts

Attention enrichment is the $\log_2$ ratio of a lineage's attention share to its cell fraction; its association with age is tested by partial Spearman correlation adjusted for log lineage fraction. Readouts and replacements reweight each sample to the mean CIMA lineage composition, renormalized over the lineages present. Shapley values are computed exactly over all coalitions of the six lineages. Equal-count replacement exchanges, in each recipient, $k$ cells (the smallest of its B, CD4 T, CD8 T and NK counts) of one lineage, or $k$ random cells as control, with same-lineage cells of a donor; changes are averaged over 50 draws and contrasted between the two exchange directions of each pair. Programme scores are means of $\log(1 + c_{\mathrm{norm}})$ over the genes in Supplementary Table 3, with $c_{\mathrm{norm}}$ counts per ten thousand, averaged within lineage and participant. CIMA-trained ridge readouts predict the CD4 cytotoxic and B secretory programmes, with regularization selected by participant-fold validation error and training-fold centring and scaling retained for external cohorts. The B-local readout fixes non-B lineages to a common background. Treatment programme changes back-project RNA changes through training loadings.

#### B-cell states and RNA decomposition

Immunobiology B-cell labels are transferred to Hashimoto B cells by participant- and class-balanced ridge classification on UCE features. Cells are called plasma-like when the plasma score exceeds the highest other score by a margin set for out-of-fold precision of at least 0.90; other cells take the higher of the naive and memory scores. The sensitivity analysis excludes 0%, 10% or 20% of cells with the smallest top-two margin. With $p_{is}$ the fraction of participant $i$'s B cells in state $s$, $c_{is}$ the programme sum in state $s$ divided by the B-cell count, participant-equal group means $p_{gs}$ and $c_{gs}$, and $\mu_{gs} = c_{gs}/p_{gs}$,

$$\Delta_{\text{programme}} = \sum_s \Big[\underbrace{(p_{\text{SC},s} - p_{\text{CT},s})\tfrac{\mu_{\text{SC},s}+\mu_{\text{CT},s}}{2}}_{\text{allocation}} + \underbrace{(\mu_{\text{SC},s} - \mu_{\text{CT},s})\tfrac{p_{\text{SC},s}+p_{\text{CT},s}}{2}}_{\text{expression}}\Big]. \tag{S9}$$

#### Statistical inference

Participants are the independent units of inference. Confidence intervals are 95% percentile intervals, paired comparisons share resampling indices, and Benjamini–Hochberg correction is applied within the families below.

| Analysis | Resampling unit and number | Test and number | Multiplicity family |
|---|---|---|---|
| Age and pooling | Participant; 10,000 bootstraps | Paired difference interval | None |
| Vaccination benchmarks | Participant; 10,000 bootstraps | 10,000 two-sided sign flips | Per task and metric: 11 error or 9 cosine contrasts |
| History substitution | Participant; 10,000 bootstraps | 100,000 two-sided sign flips | 4 cosine; separately 8 RMSE/MAE contrasts |
| Reference composition | Participant; 10,000 bootstraps | 10,000 two-sided sign flips | 12 prespecified reference-composition contrasts |
| Similar current states | Participant reference labels | 5,000 one-sided permutations | Single pairwise-cosine test |
| Response terciles | Participant; 2,000 bootstraps | Percentile interval | None |
| Window reference return | Participant; 2,000 bootstraps | 2,000 two-sided sign flips | 21 window-by-modality contrasts |
| Protein half-return | Participant paired errors | 20,000 two-sided sign flips | 1,229 proteins |
| Attention–age association | Within-cohort age labels | 10,000 permutations | Cohort-by-lineage comparisons |
| Longevity replacement | Participant; 50 cell-exchange draws averaged | All 792 group-label allocations | Prespecified lineage contrasts |
| RNA decomposition | Participant within group; 10,000 bootstraps | Percentile interval | None |
| Treatment forecasting | Patient; 10,000 bootstraps | Paired difference interval | None |
| KYDAR response | Patient, stratified by response; 10,000 bootstraps | Paired difference interval | None |

### Supplementary Tables

The accompanying workbook, `Supplementary_Tables.xlsx`, provides Supplementary Tables 1–5, one per worksheet.

#### Supplementary Table 1. Cohorts and data sources

The workbook lists cohort names, accession identifiers or source locations, platform, analysed participant and sample counts, visits, cells per sample, age ranges and use in this study.

#### Supplementary Table 2. Treatment participants and visits

The myeloma treatment subtable (GSE309595) gives patient identifiers, regimen, response grade, days from treatment start, selected cells at each of the three visits and complete-blood-count availability. A separate KYDAR subtable gives baseline identifiers, response groups, selected cell counts and sample accessions. The patient-level records are provided in the accompanying workbook.

#### Supplementary Table 3. RNA programmes

Programme scores use the mean log-transformed counts per ten thousand over the genes below.

| Programme | Genes |
|---|---|
| B-cell secretory | MZB1, JCHAIN, XBP1, PRDM1, SDC1, CD38, IGHG1 |
| Interferon response | IFIT1, IFIT2, IFIT3, ISG15, MX1, OAS1, OAS2, IRF7 |
| Cytotoxic lymphocyte | NKG7, GNLY, GZMB, GZMH, PRF1, CTSW |
| Inflammatory myeloid | IL1B, S100A8, S100A9, CTSD, FCN1, LILRB1 |
| B cell | MS4A1, CD79A, CD37, CD74, HLA-DRA, CD22 |
| Cell cycle | MKI67, TOP2A, TYMS, UBE2C, CENPF, STMN1 |

#### Supplementary Table 4. Reference-return windows

Rows follow the window order in Fig. 4D. Median intervals are calculated from actual visit dates; counts give participants with complete measurements in the cross-fitted comparisons in Fig. 4E.

| Start visit | End visit | Vacci-nated | Median days | Cell subsets | Cell types | Proteins |
|---|---|---|---|---|---|---|
| Flu Year 2 Day 0 | Flu Year 2 Day 7 | Yes | 7 | 84 | 84 | 83 |
| Flu Year 2 Day 7 | Flu Year 2 Day 90 | Yes | 84 | 82 | 82 | 81 |
| Flu Year 2 Day 0 | Flu Year 2 Day 90 | Yes | 91 | 82 | 82 | 82 |
| Immune Variation Day 0 | Immune Variation Day 7 | No | 7 | 67 | 67 | 67 |
| Immune Variation Day 90 | Flu Year 2 Day 0 | No | 70 | 64 | 64 | 64 |
| Immune Variation Day 0 | Immune Variation Day 90 | No | 93 | 65 | 65 | 65 |
| Immune Variation Day 0 | Flu Year 2 Day 0 | No | 171 | 65 | 65 | 65 |

#### Supplementary Table 5. Comparator implementations

The workbook lists the method, task, input representation, weights or training data, task adaptation, settings and model-selection procedure, and code repository and version. Vaccine entries cover the cross-year and within-season benchmarks.

#### Supplementary Table 6. Hyperparameters

| Parameter | Setting |
|---|---|
| **Architecture** | |
| UCE / projection / attention hidden dimensions | 1,280 / 128 / 64 |
| RNA dimension / slots | 16 / 8 |
| Mean / covariance basis ranks | 7 / 11 |
| Moment encoding / phase embedding / transition hidden dimensions | 64 / 16 / 128 |
| Refinement layers / width / operator rank | 4 / 64 / 8 |
| History transition / projection dimensions | 48 / 64 |
| Transformer layers / attention heads | 1 / 4 |
| Low-rank $r$ / scale $\alpha$ | 8 / 0.5 |
| KYDAR radial-basis anchors | 32 |
| **Personal reference and scaling** | |
| CLR dimension $D$ / pseudocount $\varepsilon$ | 21 / 0.5 |
| Shrinkage $\lambda$ grid | $\{0.25, 0.5, 1, 2, 4, 8, 16, 32, 64, \infty\}$ |
| Inner participant folds for reference fitting | 4 |
| Response-scale $\gamma$ interval | $[0, 1.25]$; treatment: fixed at 1 |
| Covariance regularizer $\epsilon$ | $10^{-4}\max\{\mathrm{tr}(\mathbf{C})/d, 10^{-4}\}$ |
| Eigenvalue-bound quantiles $q_{\mathrm{lo}}/q_{\mathrm{hi}}$ | 0.01 / 0.99 |
| **Optimization** | |
| Outer folds: CIMA, Sound Life, treatment, KYDAR | 5 each |
| Inner folds: Sound Life / treatment / KYDAR | 4 / 3 / 3 |
| Optimizer / weight decay for neural stages | AdamW / $10^{-4}$ |
| CIMA age: learning rate; participants per update; maximum / minimum epochs; patience; gradient clip | $10^{-3}$; 8; 300 / 15; 20; 1 |
| Sound Life and treatment affine: learning rate; participants per update; maximum epochs; patience; gradient clip | $10^{-3}$; 16; 300; 25; 5 |
| Sound Life and treatment refinement: learning rate; participants per update; maximum epochs; patience; gradient clip | $3 \times 10^{-4}$; 8; 150; 15; 5 |
| Sound Life history: learning rate; participants per update | $10^{-3}$; 8 |
| Treatment history: learning rates; patients per update; maximum epochs; patience | $\{3 \times 10^{-4}, 10^{-3}\}$; 8; 25; 12 |
| KYDAR affine: learning rate; maximum epochs; patience | $10^{-3}$; 150; 20 |
| KYDAR refinement: learning rate; maximum epochs; patience | $3 \times 10^{-4}$; 80; 12 |
| KYDAR head: inverse regularization; state-feature multiplier; Brier penalty | $\{0.01, 0.1, 1, 10\}$; $\{0.25, 1\}$; 0.2 |
| UCE cells per query / support visit, Sound Life | 512 / 512 |
| RNA cells: affine / refinement training, Sound Life | 2,048 / 512 |
| Virtual cells per prediction, Sound Life | 2,048 |
| Age UCE cells: CIMA / Immunobiology / OneK1K | 1,024 / 1,500 / up to 1,500 |
| Hashimoto cells per primary sample | 1,500 |
| Treatment / KYDAR cells per visit | 512 / 64 |
| History temperatures $\tau_1/\tau_2$ | 0.05 / 0.1 |
| History contrast / current-anchor weights | 0.5 / 0.25 |
| Geometry vector / Gram weights; scale floor | 0.75 / 0.25; $10^{-3}$ |
